# Investigating the Effects of Psilocybin on Cognitive Flexibility in Touchscreen and Naturalistic Variations of the Probabilistic Reversal Learning Task

**DOI:** 10.64898/2026.08.06.743309

**Authors:** Dasha Anderson, Nanni Maillot, Christopher W Thomas, Caroline T Golden, Gary Gilmour, Emma SJ Robinson

## Abstract

**Rationale:** Psychedelic compounds such as psilocybin have attracted growing interest for their potential therapeutic effects in psychiatric disorders, with improvements in cognitive flexibility proposed as a possible mechanism of action. However, the effects of psychedelics on cognitive flexibility remain poorly understood.

**Objective:** This study aimed to examine the acute and post-acute effects of psilocybin (0.1, 0.3, 1 mg/kg) and lysergic acid diethylamide (LSD, (0.02, 0.04, 0.08 mg/kg) on cognitive flexibility in male rats.

**Methods:** This was tested using two variants of the probabilistic reversal learning task (PRLT): a touchscreen-based operant task and a more ethological foraging-based task.

**Results:** In the touchscreen PRLT, acute psilocybin disrupted task engagement, with animals completing fewer trials and showing increased trial initiation latency, although psilocybin also showed a trend toward faster initial rule acquisition. However, psilocybin did not significantly alter the number of rule changes achieved, a canonical measure of cognitive flexibility, or feedback sensitivity. LSD similarly produced limited acute effects, although the highest dose reduced lose-shift probability, suggesting decreased sensitivity to negative feedback under some conditions. Post-acute effects of psilocybin were minimal in both PRLT variants and, where LSD effects were observed these occurred across different doses and timepoints without a consistent pattern.

**Conclusions:** Overall, these findings suggest that serotonergic psychedelics do not robustly enhance reversal learning in these paradigms and that apparent learning effects may reflect transient disruptions in task engagement rather than improvements in cognitive flexibility. These results also highlight potential limitations of these PRLT paradigms for detecting psychedelic-induced changes in cognitive flexibility in rodents.

## Introduction

Psychedelics are gaining attention for their potential utility in the treatment of psychiatric conditions with psilocybin achieving rapid-acting antidepressant effects in clinical trials for treatment-resistant depression (Carhart-Harris et al., 2016; Goodwin et al., 2022; Raison et al., 2023; Von Rotz et al., 2023). Despite this, the precise mechanisms underlying the therapeutic effects of psychedelics remain unclear. In humans, psychedelics appear to improve cognitive flexibility (Doss et al., 2021; Davis et al., 2020) – the ability to change one’s cognitive state to effectively adapt to environmental changes (Johnco et al., 2014). Assessing these domains in rodents provides a possible translational method to investigate the mechanisms underlying psychedelic treatment effects. There are two primary methods for assessing cognitive flexibility: set-shifting tasks, which involve shifting attention between different stimulus dimensions (e.g. texture vs odour), and reversal learning tasks (RLTs), which test the ability to update choice behaviour based on changing reward contingencies (Anderson and Robinson, 2025).

In RLTs, reward contingencies can either be deterministic or probabilistic. 5-HT2A receptor agonists have shown mixed effects in cognitive flexibility tasks in rodents (Amodeo et al., 2020; Torrado Pacheco et al., 2023; Odland et al., 2021; Conn et al., 2024; Šabanović et al., 2024; Fisher et al., 2024). So far, psilocybin has not been tested in a PRLT, a paradigm well-suited to probing cognitive flexibility under uncertainty which arguably closer represents the ambiguity of real-life decision-making. Subjects are presented with a choice between two stimuli – each associated with different probabilities of receiving a reward. Over time, the reward contingencies associated with the stimuli are reversed, such that the previously rewarded stimulus becomes unrewarded and vice versa, challenging subjects to flexibly adjust their behaviour based on probabilistic feedback. This enables measurement of several behavioural domains, including cognitive flexibility, task engagement and performance, and reward sensitivity. The PRLT can also be combined with computational reinforcement learning models to examine computational processes underlying choice behaviour based on trial-by-trial behaviour (Izquierdo et al., 2017). Importantly, this task has a direct equivalent that can be tested in human populations, allowing back-translation. Indeed, depressed patients show impairments in this task (Mukherjee et al., 2020).

Here, we aimed to investigate dose-dependent acute and sustained effects of psilocybin and a comparator, LSD, in two variants of the PRLT in rats: a touchscreen (Bari et al., 2010; Wilkinson et al., 2020) and a foraging (bowl-digging) variant (Jackson et al., 2024; Griesius et al., 2023). We included both versions of the task as the touchscreen method requires prolonged training schedules biasing behaviour towards procedural learning (Jackson et al., 2024) while the foraging task requires minimal training and favours more naturalistic and dynamic behaviour (Jackson et al., 2024). If psychedelic treatment increases cognitive flexibility in these tasks, an increase in the number of rule changes achieved in a session is predicted.

## Methods

### Animals

Two cohorts of 12 male Lister-Hooded rats (Envigo, UK) (total 24 animals, 1 animal = 1 experimental unit) were pair-housed in enriched housing containing a red shelf, wooden chews, rope, and cardboard tube. Rats were provided with access to a playpen for up to 30 min per day during weeks when they were not training or testing (www.3hs-initiative.co.uk). Rats underwent 5 days of habituation to handling prior to starting behavioural training. The sample size was based on previous rodent PRLT studies which we expected elicited a similar effect size to the current study (Jackson et al., 2024; Wilkinson et al., 2020). Animals were kept in temperature- (21±1°C) and humidity- (45-65%) controlled conditions under a 12:12 reverse light-dark cycle (lights off at 8.15am). Animals were kept on a restricted diet of 18g laboratory chow (LabDiet, USA) per rat but maintained at no less than 90% their free-feeding weight. All studies were performed in accordance with with the UK Animal (Scientific Procedures) Act, with ethical approval from the University of Bristol AWERB and under a Home Office Project license (PPL number PP4130557). The touchscreen cohort weighed 465±8g before training, 511±10g before the drug studies, and 543±39g after drug studies. The bowl digging cohort weighed 465±8g before training, 511±10g before the drug studies, and 543±39g after the drug studies. All behavioural experiments were conducted during the dark phase.

### Drugs

COMP360 (Compass Pathways’ proprietary formulation of psilocybin, from now ‘psilocybin’) was provided by Compass Pathways (UK) and LSD was obtained from Cambridge Biosciences. Compounds were dissolved in sterile saline before use (pH=7). Psilocybin was administered intraperitoneally (i.p.) using a refined handling method (Stuart et al., 2015, www.3hs-initiative.co.uk) at 0 (vehicle control), 0.1, 0.3, and 1 mg/kg in the touchscreen PRLT, however 0.1 mg/kg was not included in the bowl digging PRLT due to the lack of effects at this dose in the touchscreen task. LSD was administered at the following doses: 0, 0.02, 0.04, 0.08 mg/kg (also i.p.) and only tested in the touchscreen task. Aliquots were shielded from light with foil during use. Doses were selected with the aim of eliciting clinically relevant exposure based on previous work from our lab (Hinchcliffe et al., 2024).

### Touchscreen training

Rats were tested in sound-proofed operant boxes containing a three-panel infrared touchscreen (Med Associates Inc, USA), house light and tone generator (**Fig 1A**) as previously described in Jackson et al. (2024) and Wilkinson et al. (2020). Rats could collect 45mg reward pellets (LabDiet, UK) from a magazine opposite the touchscreen. Stimuli were controlled and responses recorded via KLimbic software (Med Associates Inc, USA). Training began with continuous reinforcement training (CRF) in which an animal learned to touch an initiation square in the central window to obtain a reward pellet. The session ended after 30 minutes or completion of 120 trials. To pass onto the next stage, rats needed to complete at least 120 trials over two consecutive days. Animals then learned to touch the initiation square, followed by either the left or right window to obtain a reward pellet. The session ended after 20 minutes or completion of 200 trials. If the rat touched the initiation square but failed to touch either window within 10 seconds, a 10 second time-out period occurred where the house light turned on and no reward pellet was delivered. Training data are presented in **Supplementary Fig S1**.

**Fig 1.**
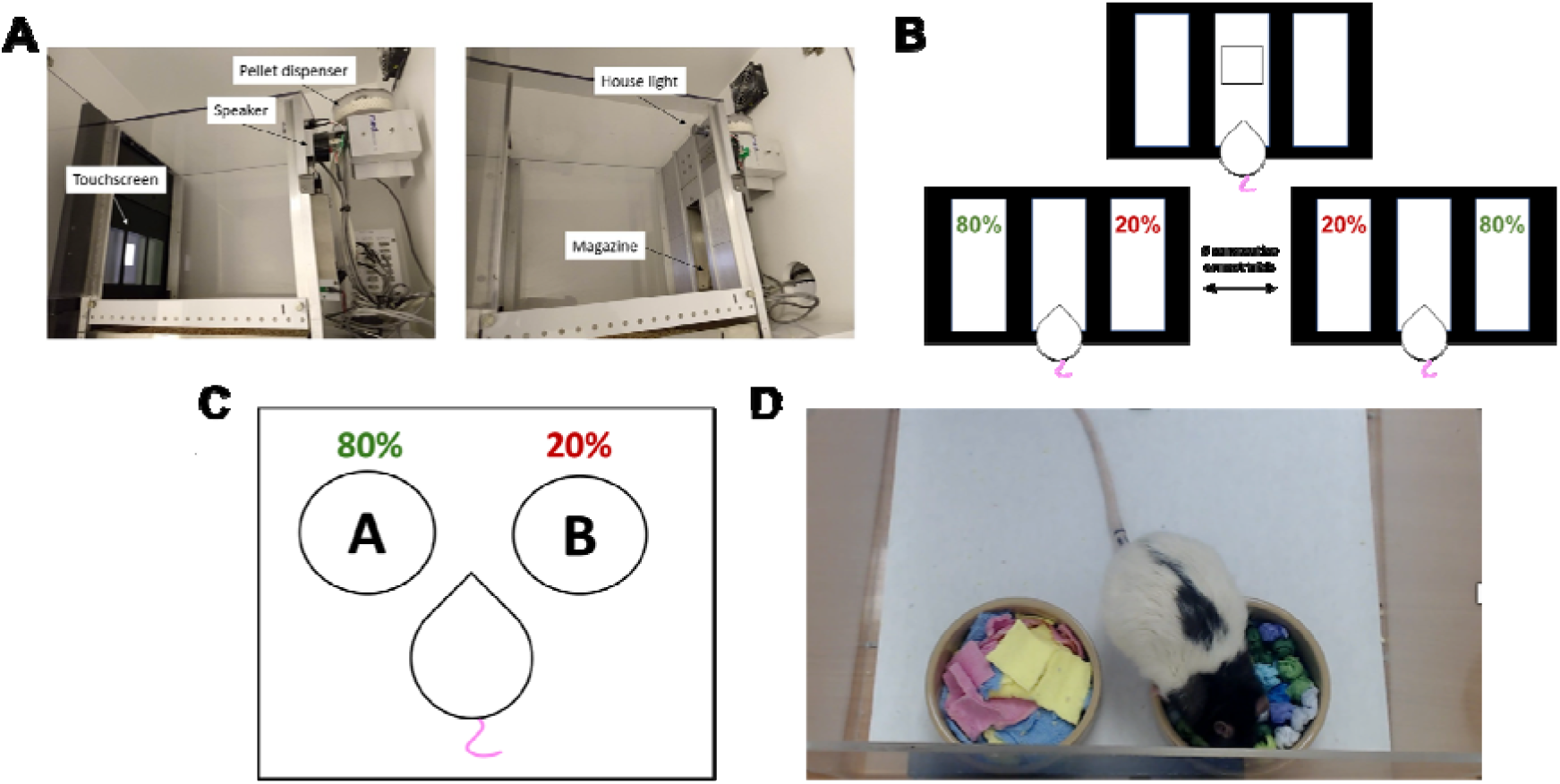
Probabilistic Reversal Learning Task (PRLT). (**A**) Operant system apparatus, including view from right side of operant chamber (left) and view from left side of operant chamber (right). (**B**) Trial structure of the touchscreen PRLT. The rat presses one of two touchscreens, one of which is rewarded 80% of the time and the other 20% of the time. (**C**) Trial structure of the foraging PRLT. The rat chooses to dig in one of two bowls, containing two different digging materials, one of which is rewarded 80% of the time and the other 20% of the time. (**D**) Example image of rat digging in materials in the experimental set up.

### Touchscreen Probabilistic Reversal Learning Task

After reaching at least 120 trials over two consecutive sessions in CRF2, they moved onto the PRLT (Bari et al., 2010; Wilkinson et al., 2020). Either the left or right window was rewarded 80% of the time (the ‘rich’ stimulus) and the other 20% of the time (the ‘lean’ stimulus) (**Fig 1B**). The starting position of the rich stimulus was counterbalanced across rats and remained the same across sessions. After rats selected the rich stimulus on 8 consecutive trials, the reward contingencies switched so that the previously rich stimulus became the lean stimulus and vice versa (“rule change”). This continued every eight consecutive rich choices until animals completed 200 trials or 40 minutes passed. Unrewarded stimuli and omissions (no stimulus choice within 10 seconds) were followed by a 5 second time-out period where the house light came on. Baseline data are presented in **Fig S2**.

Raw and Q-learning output measures were calculated using Matlab 2022a (Mathworks, USA). Task performance and engagement were assessed by the total number of trials completed, trial initiation time, and accuracy (percentage of rewarded trials). The number of trials until the first rule change was achieved provided a measure of reward learning and the overall number of rule changes provided a measure of cognitive flexibility. Two feedback sensitivity measures were also calculated: rewarded outcomes followed by the decision to stay with the same stimulus (win stay/positive feedback sensitivity) and unrewarded outcomes followed by the decision to change stimulus (lose shift/negative feedback sensitivity). Feedback could further be classified as “true” or “misleading”, depending on whether the feedback matched the usual feedback for that stimulus.

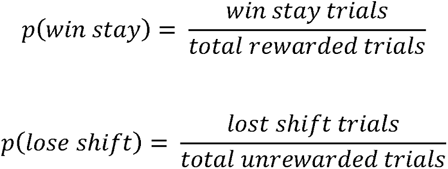

In addition to raw output measures, two computational reinforcement learning models were fit to the animals’ trial-by-trial PRLT data. Full modelling methods are provided in **Supplementary Methods S3** but, in brief, the first model (Q-learn1) provided two parameters additional to the raw behavioural measures described above: learning rate (*α*) and choice determinism (*β*). In the second model (Q-learn2), learning rate was split by positive and negative feedback, resulting in three additional parameters: positive learning rate, negative learning rate, and *β* value.

### Foraging training

Similar training methods were used as described in Jackson et al. (2024). Full foraging study methods are provided in **Supplementary Materials S4**. Animals were habituated to a 40 cm^2^ Perspex arena with a tray liner and two ceramic bowls placed at the back of the arena approximately 1 cm apart (**Fig 1C**). Bowls were baited with 45mg reward pellets (LabDiet, UK) as shown in **Fig 1D**. They then completed an initial reward-retrieval session in which bowls baited at different clock-face positions contained a pellet. Animals completed 12 trials and were briefly removed if they failed to retrieve the pellet within 30 seconds. Next, rats received three days of digging training. Pellets were placed in a fixed location but buried under increasing depths of sawdust, with 12 trials per session. Failure to find the pellet within 15 seconds was scored as an omission. Following training, rats completed three discrimination tests using different digging substrates. One substrate was consistently baited and counterbalanced across animals, with crushed pellet added to both substrates to control for olfactory cues. Animals were considered fully trained after six consecutive correct choices within 20 trials. Discrimination training data are presented in **Supplementary Fig S5**. Finally, rats performed a probabilistic learning session in which one substrate was rewarded on 80% of trials (rich) and the other on 20% (lean). Animals had to make six consecutive correct choices within 30 trials to acquire the rule, with omissions recorded if no choice was made within 30 seconds.

### Foraging Probabilistic Reversal Learning Task

Following successful probabilistic learning (without reversal), animals underwent a baseline PRLT session. Rats were split into two groups of six and run in staggered sessions. The rat had to choose the rich substrate 6 times consecutively within a maximum of 50 trials to successfully acquire the first rule. Once the rat had acquired this initial rule, the contingencies were reversed such that the rich substrate now became the lean substrate and vice versa. The reward contingencies were then reversed every time the rat chose the rich bowl 6 times consecutively up to a maximum of 7 reversals. The position of the bowl was pseudo-randomised across trials and the substrate to be rewarded first was counterbalanced across animals. A 30 second cut-off was used again. The session ended after one hour, five consecutive omissions, or after failing to acquire a rule within 50 trials. Similar raw output measures were obtained to the touchscreen PRLT.

### Experimental schedule

In acute touchscreen PRLT studies, animals were dosed and tested twice a week. Animals were dosed and then tested one hour later.. In post-acute studies, animals were dosed once a week and tested at two timepoints after dosing (24 hour and 7 days). Acute studies were fully counterbalanced to reduce the impact of any carry over effects. Given evidence of acute psilocybin effects on motivation and engagement in the touchscreen PRLT, only post-acute timepoints were used for the foraging PRLT study: 24 hours and 6 days. Animals were dosed once a week and tested 24 hours and 6 days after dosing in a staggered manner. Due to the repeated measures design and the length of PRLT sessions, it was not feasible to include a 7-day timepoint, so we instead opted for a 6-day timepoint. All animals underwent all drug conditions and timepoints in a randomised order in a within-subjects fully counterbalanced design, with experimenters blinded to treatment group throughout the entire study. Separate cohorts were used for touchscreen and foraging experiments. An *a priori* criterion was set where if animals completed less than 50 out of 200 trials in one touchscreen PRLT session, they were excluded from analysis of that session and their data replaced with the group mean. Such low levels of task completion are uncommon and are generally indicative of unrelated factors, such as technical issues affecting task delivery. One animal performed <50 trials in over half the sessions and was thus excluded from analysis, as its behavioural data could not be considered a reliable measure of PRLT performance.

### Statistical analysis

Statistics were performed in SPSS version 30.0.0.0 (IBM, USA) and GraphPad Prism 9.0.0 (GraphPad Software, USA). Prism was also used to create figures. Outliers outside two standard deviations of the group mean were replaced with the group mean, which was decided *a priori*. Data were normality tested and two-tailed one-way repeated measures ANOVAs or appropriate non-parametric equivalents (e.g. Friedman’s test) were used to assess acute psilocybin effects on PRLT measures. Significant main effects were followed up with Dunnett’s post hoc tests comparing each drug condition to vehicle. Two-way repeated measures ANOVAs were used to assess post-acute effects of psilocybin on all PRLT measures, with dose and time included as fixed factors. Significant interactions were further explored by performing Dunnett’s tests at each timepoint. Only significant main effects of dose were followed up with post hoc Dunnett’s tests based on estimated marginal means, where interactions were not significant. Trend level main effects (p<0.1) are reported but not further analysed. All data were sphericity tested and the Huynh-Feldt correction applied if Mauchley’s test of sphericity was violated. Statistical trends were defined as *p* values between *p*=0.05 and *p*=0.1. Data are presented as mean ± SEM. Pairwise comparisons vs vehicle control are plotted on figures, *≤0.05, **<0.01, ***<0.001.

## Results

### Acute psilocybin effects on the touchscreen PRLT

Psilocybin showed some evidence of acute effects on learning in this task, significantly reducing the number of trials to the first rule change (F(3, 33)=3.242, *p*=0.034, n=12) (**Fig 2A**). There was a trend towards rats learning the first rule in significantly fewer trials following 1 mg/kg psilocybin (*p*=0.061). While psilocybin appeared to affect the number of rule changes achieved in a session (F(3, 33)=3.585, *p*=0.024, n=12), with pairwise comparisons revealing a significant reduction at 1 mg/kg (*p*=0.034) (**Fig 2B**), however this was no longer significant when was normalised to trials completed (F(3, 33)=0.308, *p*=0.819, n=12). Regarding feedback sensitivity, acute psilocybin had no significant effect on win stay probability (F(3, 33)=1.65, *p*=0.191, n=12) (**Fig 2C**), however there was a trend toward a significant effect of acute psilocybin on overall lose shift probability (F(3, 33)=2.816, *p*=0.054, n=12) (**Fig 2D**). No clear effects of acute psilocybin were found on these measures when split by positive or negative feedback (**Supplementary Fig S6**). Psilocybin showed acute effects on Q-learning outputs, increasing overall Q-learn1 learning rates (F(3, 33)=9.085, *p*<0.001, n=12), and this appeared to be specific to negative learning rates when *α* values were split by positive and negative feedback in Q-learn2 (X^2^(4)=14.10, *p*=0.003, n=12) (**Supplementary Fig S7A,D**). However, the latter model showed poorer fit to the data based on Bayesian Information Criteria (BIC) values (*t*=9.973, df=11, *p*<0.001, n=12). Notably, psilocybin also showed non-specific effects on task engagement and performance, significantly affecting the number of trials completed in a session (X^2^(4)=23.26, *p*<0.001, n=12), which were reduced at 1 mg/kg (*p*<0.001) (**Fig 2E**), as well as trial initiation time (F(3, 33)=16.97, *p*<0.001, n=12), which was increased at 1 mg/kg (*p*=0.001) (**Fig 2F**). Acute psilocybin also induced a significant main effect on accuracy (F(3, 33)=3.975, *p*=0.016, n=12), however post hoc tests were not significant (*p*≥0.099) (**Fig 2G**).

**Fig 2.**
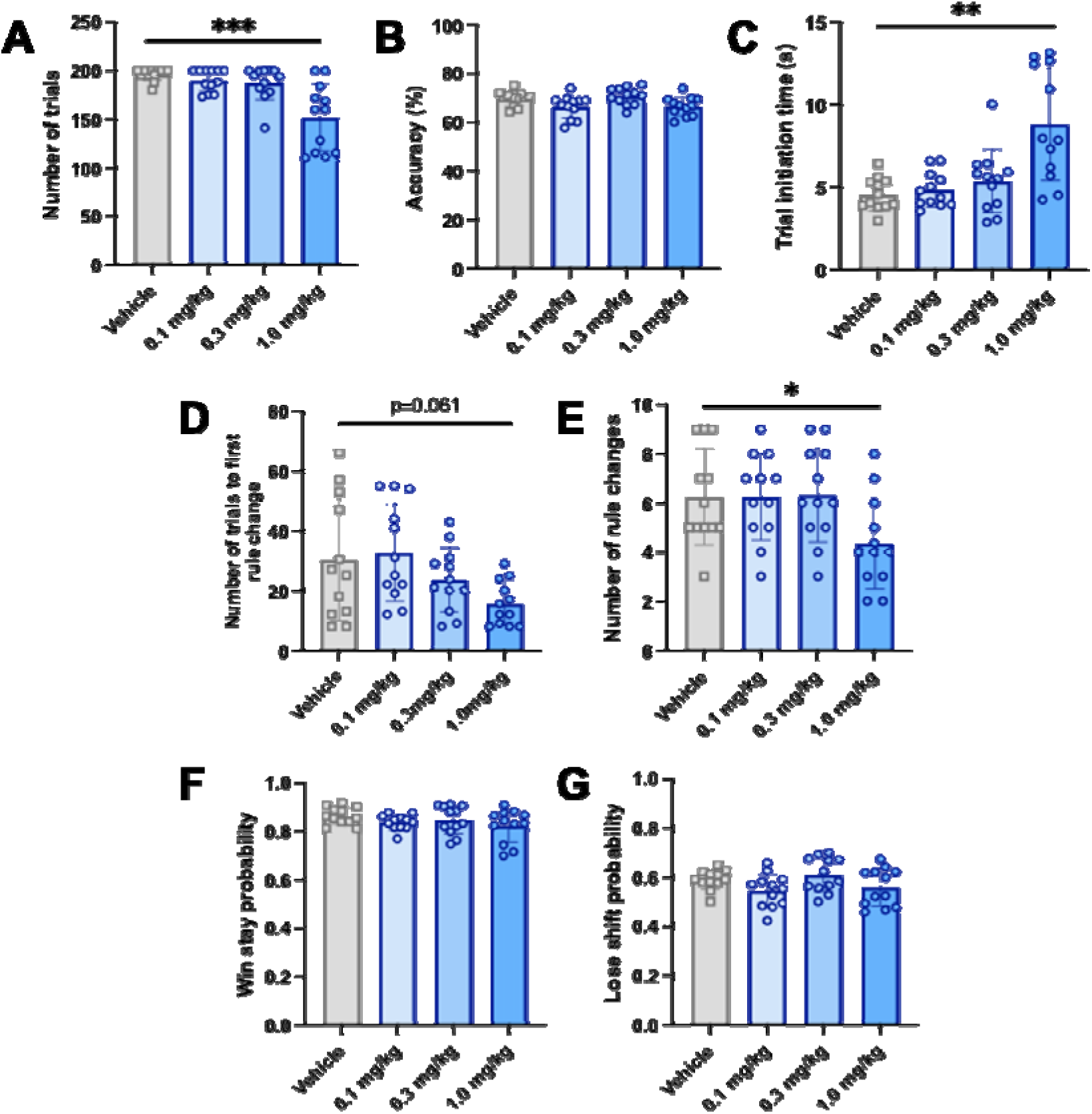
Acute effects of psilocybin on main operant PRLT output measures. (**A**) Number of trials completed in a session. (**B**) Accuracy (percentage of total trials that were correct responses). (**C**) Time taken for animals to self-initiate each trial. (**D**) Number of rule changes completed in a session. (**E**) Trial at which animals achieved their first rule change in a session. (**F**) Win stay probability (likelihood of selecting the previously rewarded stimulus). (**G**) Lose shift probability (likelihood of avoiding the previously rewarded stimulus). Data shown as mean +/- sem and individual data points represent individual subjects, n = 12, *p<0.05, **p<0.01, ***p<0.001 post-hoc pairwise comparison vs vehicle control

### Post-acute effects of psilocybin on the touchscreen PRLT

Psilocybin showed negligible post-acute effects when animals were tested at 24hrs and 7 days post-treatment. There was a trend interaction between psilocybin dose and time on the number of trials to the first rule change (F(3, 33)=2.416, *p*=0.084, n=12) (**Fig 3A**) and while there was a significant main effect of time on rule changes (F(1, 11)=24.87, *p*<0.001, n=12), there was no significant effect of psilocybin dose (F(3, 33)=1.785, *p*=0.169, n=12) (**Fig 3B**). No significant main effect of dose, time or interaction between these factors was found on overall win stay or lose shift probability (*p*≤0.277) (**Fig 3C**), while minimal effects of acute psilocybin were shown on feedback sensitivity measures when split by feedback type (**Supplementary Fig S8**). No clear post-acute effects of psilocybin were shown on any Q-learning model parameters (**Supplementary Fig 39**). There was some evidence of post-acute psilocybin effects on task performance, with significant main effects of dose (F(3, 33)=4.674, *p*<0.001, n=12) and time (F(1, 11)=8.799, *p*=0.013, n=12) on trials completed and a trend toward an interaction between these factors (F(3, 33)=2.666, *p*=0.064, n=12) (**Fig 3E**). Post hoc tests showed trends towards completion of more trials following 0.1 mg/kg (*p*=0.067) and 0.3 mg/kg psilocybin (*p*=0.086) (**Fig 3F**). While there were no post-acute effects on accuracy (*p*≥0.299), there was a significant main effect of time (F(1, 11)=17.643, *p*<0.001, n=12) but only a trend effect of psilocybin dose on trial initiation times post-acutely (F(3, 33)=2.577, *p*=0.070, n=12) (**Fig 3G**).

**Fig 3.**
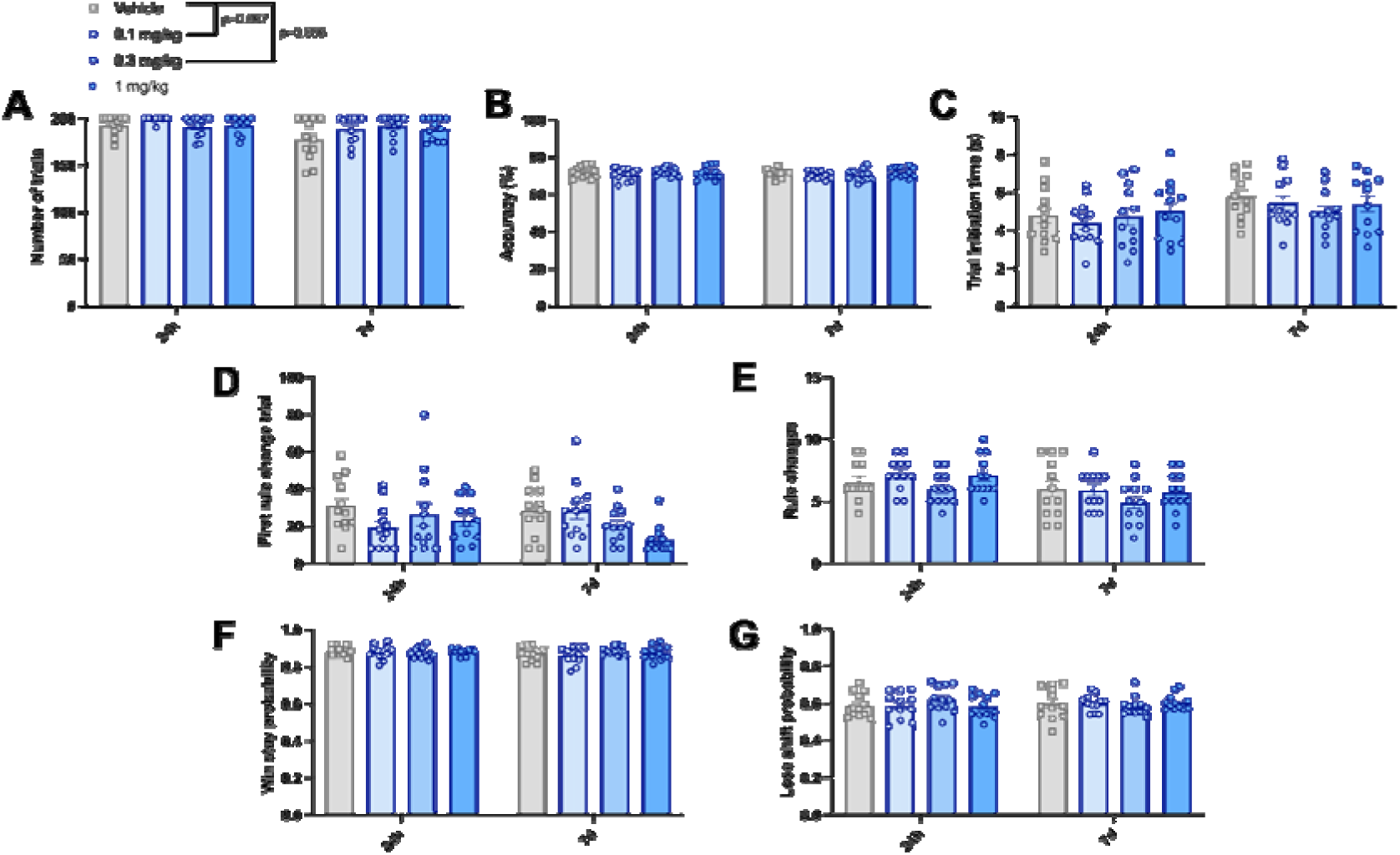
Post-acute effects of psilocybin on main operant PRLT output measures at 24 hours and 7 days post-dose. (**A**) Number of trials completed in a session. (**B**) Accuracy (percentage of total trials that were correct responses). (**C**) Time taken for animals to self-initiate each trial. (**D**) Number of rule changes completed in a session. (**E**) Trial at which animals achieved their first rule change in a session. (**F**) Win stay probability (likelihood of selecting the previously rewarded stimulus). (**G**) Lose shift probability (likelihood of avoiding the previously rewarded stimulus). Data shown as mean +/- sem and individual data points representing each subject, n = 12, *p<0.05, **p<0.01, ***p<0.001 post-hoc pairwise comparison vs vehicle control

### LSD effects on the touchscreen PRLT

LSD had limited acute effects on the main touchscreen PRLT measures (**Supplementary Fig S10**), except for producing a significant reduction in lose shift probability (F(3, 33)=3.167, *p*=0.012, n=11) at the highest 0.08 mg/kg dose (*p*=0.012) (**Supplementary Fig S10D**). When split by positive and negative feedback, this effect was only seen for lean lose shift probability (F(3, 33)=4.457, *p*=0.010, n=11) and at the highest dose (*p*=0.010) (**Supplementary Fig S11D**). LSD showed more heterogenous post-acute effects on touchscreen PRLT performance. There was a dose-by-time interaction on the first rule change trial (F(3, 33)=3.605, *p*=0.025, n=11), with animals taking significantly longer to learn the first rule 24 hours after 0.04 mg/kg LSD (*p*=0.048) (**Supplementary Fig S13D**). There was a significant post-acute effect of LSD dose on rich win stay probability (F(3, 30)=1.707, *p*=0.021, n=11), whereby rich win stay probability was significantly lower after 0.02 mg/kg LSD (*p*=0.131) (**Supplementary Fig S14B**). LSD also showed some post-acute effects on performance and motivation. There was a significant interaction between dose and time on the number of trials animals completed in a session (F(3, 30)=4.559, *p*=0.001, n=11), with animals completing significantly fewer trials 7 days after receiving 0.02 mg/kg LSD (*p*<0.001) (**Supplementary Fig S13A**). There was a significant effect of dose on trial initiation time (F3, 30)=2.958, *p*=0.048, n=11), where animals took significantly longer to initiate trials following 0.04 mg/kg (*p*=0.012) and 0.08 mg/kg (*p*=0.030) (**Supplementary Fig S13C**). The only Q-learn1 parameter post-acutely affected by LSD was β, which showed a significant interaction between dose and time (F(3, 30)=2.964, n=11), *p*=0.048), with post hoc tests revealing a trend toward reduced beta values (*p*=0.081) (**Supplementary Fig S15B**). The only Q-learn2 parameter significantly affected by post-acute LSD dose was positive learning rates (F(3, 30)=2.968, *p*=0.048, n=11), which were significantly lower following 0.02 mg/kg LSD (*p*=0.021) (**Supplementary Fig S15C**). One rat was excluded from the post-acute LSD study as he stopped performing the task reliably, achieving less than the minimum number of trials in two out of four sessions.

### Post-acute effects of psilocybin in the foraging PRLT

#### Drug study

Psilocybin treatment had limited post-acute effects when tested in a foraging version of the PRLT. There was only a trend toward a main effect of psilocybin on the number of trials required to reach the first rule change (F(1, 11)=9.186, *p*=0.092, n=12) and a significant main effect of time (F(1, 11)=9.186, *p*=0.011, n=12) (**Fig 4D**). There were trend effects of time for overall lose shift probability (F(1, 11)=4.266, *p*=0.063, n=12) (**Fig 4G**) and rich lose shift probability (F(1, 11)=3.679, *p*=0.081, n=12) (**Supplementary Fig S16B**). There was also a significant main effect of dose on rich win stay probability (F(2, 22)=4.058, *p*=0.032, n=12) (**Supplementary Fig S16A**), however post hoc comparisons were not significant (*p*≥0.238). There was a significant dose x time interaction on the number of omissions (F(2, 22)=4.791, *p*=0.019, n=12), with trends towards fewer omissions following 1 mg/kg at 24 hours (*p*=0.063) and more omissions at this dose after 6 days (*p*=0.092) (**Fig 4C**).

**Fig 4.**
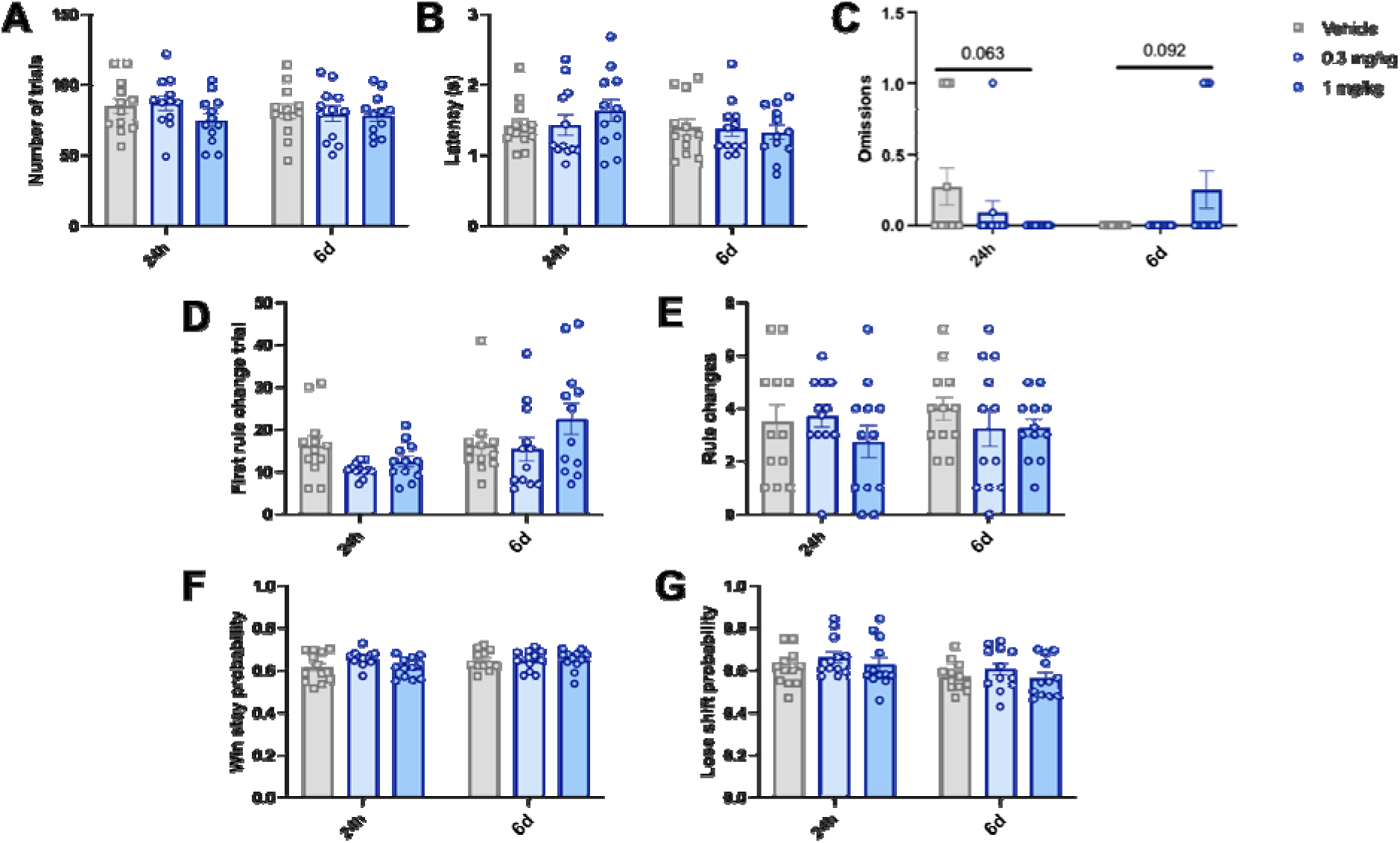
Post-acute effects of psilocybin on main foraging PRLT output measures 24 hours and 6 days post-dose. (**A**) Number of trials completed in a session. (**B**) Omissions (number of times animals took longer than 30 seconds to make a choice and the trial was restarted). (**C**) Time taken for animals to make a choice. (**D**) Trial at which animals achieved their first rule change in a session. (**E**) Number of rule changes completed in a session. (**F**) Win stay probability (likelihood of selecting the previously rewarded stimulus). (**G**) Lose shift probability (likelihood of avoiding the previously rewarded stimulus). Data shown as mean +/- sem and individual data points represent individual subjects, n = 12, *p<0.05, **p<0.01, ***p<0.001 post-hoc pairwise comparison vs vehicle control

## Discussion

Psilocybin induced mixed acute effects on cognitive flexibility measures in the touchscreen PRLT, with some evidence supporting effects on reward learning but also evidence of disruption to general task engagement. This raises the possibility that psilocybin effects on learning in this assay result from more general acute disruption. Post-acute effects of psilocybin were limited in both a standard touchscreen PRLT and a more ethologically valid foraging-based PRLT, suggesting psilocybin exhibits limited post-acute effects on cognitive flexibility in these paradigms. A similar lack of effects was seen with LSD treatment in the touchscreen task.

At 1 mg/kg, psilocybin disrupted task engagement in the touchscreen variant of the PRLT, with fewer trials completed in a session and increased trial initiation time. This indicates generalised effects on motivation, arousal and/or motor function, likely related to its perceptual effects. In line with this, reinforcement learning modelling indicated reduced overall learning rates in the touchscreen task, consistent with a general slowing of learning. Paradoxically, however, 1 mg/kg psilocybin trended toward acutely reducing the number of trials required to acquire the first rule, suggesting enhanced stimulus-outcome learning. A simple Q-learning model with a combined learning rate indicated slowed learning overall, a dual learning rate model revealed selectively reduced learning from negative RPEs. This could support reward learning by limiting over-adjustment to unrewarded trials, allowing rats to accumulate more evidence regarding the correct choice. However, BIC values indicated better fit of the simpler model to the present data than the asymmetric model and psilocybin effects on rule acquisition were only trend level.

Psilocybin lacked acute effects on more canonical measures of cognitive flexibility and reward sensitivity. Contrary to our prediction, acute psilocybin did not alter total rule changes achieved in a session. Initially, this seems inconsistent with acute psilocybin improvement of rule switching in a rodent set-shifting task (Torrado Pacheco et al., 2023). However, set-shifting and reversal learning are underpinned by distinct cognitive and neural substrates (Hamilton and Brigman, 2015; Kesner and Churchwell, 2011). Whereas reversal learning involves learning of new stimulus-outcome associations (Izquierdo et al., 2017), set-shifting requires shifting attention and behaviour between previously learned strategies (Birrell and Brown, 2000). Furthermore, Torrado Pacheco et al. (2023) found no acute effects of psilocybin on deterministic RLT, consistent with the present findings. Our findings also align with evidence that the 5HT2A/C receptor agonist DOI does not affect reversal learning in a rodent spatial PRLT (Amodeo et al., 2020). Likewise, psilocybin did not acutely modulate win stay and lose shift measures, despite evidence of heightened sensitivity to negative feedback in MDD (Tavares et al., 2008; Murphy et al., 2003) and established serotonergic modulation of these measures (Bari et al., 2010; Evers et al., 2005).

LSD similarly showed limited acute effects on learning and cognitive flexibility in the PRLT. However, at the highest dose tested (0.08 mg/kg), LSD reduced overall and lean lose-shift probability, suggesting decreased sensitivity to negative feedback under some conditions. Despite this, effects were otherwise minimal, indicating that serotonergic psychedelics do not robustly alter cognitive flexibility as measured in this task. The present findings also underscore the importance of assessing drug effects on general task performance to disentangle genuine learning improvements from transient impairments.

Psilocybin had limited post-acute effects in touchscreen PRLT measures, indicating that any apparent acute facilitation of rule learning observed was not sustained. The only post-acute changes observed were trends towards increased numbers of trials completed after 0.1 and 0.3 mg/kg although the lack of dose-dependence limits interpretation. The effect appears to be primarily driven by a decline in control group performance, especially after 7 days, rather than psilocybin-induced improvement, which may involve a buffering effect against natural drops in motivation or task persistence. However, this effect was statistically weak. The lack of post-acute improvements in cognitive flexibility seems at odds with human data (Doss et al., 2021; Murphy-Beiner and Soar, 2020) measured using set-shifting tasks, such as the Wisconsin Card Sorting Task (WCST) and the Penn Conditional Exclusion Test (PCET). LSD produced more varied post-acute effects on touchscreen PRLT performance. Some behavioural changes were observed, but these occurred at different doses and timepoints rather than following a clear dose-response pattern.

Psilocybin similarly exhibited limited effects in the foraging PRLT 24 hours and 6 days post-dose. Psilocybin had no consistent post-acute effects on general task performance measures such as trials completed and latency, indicating any acute effects reverse within 24hrs and there are no sustained beneficial or detrimental effects. There was a trend towards psilocybin-induced effects on omissions, with the high dose reducing and increasing omissions after 24 hours and 6 days, respectively. However, the 24-hour effect appeared strongly driven by increased vehicle group variance not observed at 6 days post-dose, rather than a drug-induced improvement. With most animals showing no omissions, apparent group differences predominantly reflected isolated cases (e.g. a single omission in a few animals), with a single trial change substantially altering variance. Thus, these effects are unlikely to reflect a reliable post-acute influence of psilocybin.

The foraging PRLT is more ethologically relevant than the touchscreen PRLT. While the touchscreen variant allows strong experimental control, it is highly artificial, requiring animals to learn the task over a prolonged period of training. which may result in a bias towards procedural learning. Learning in certain modalities (e.g. odour, digging) is more easily acquired in rodents than visual or touchscreen tasks (Izquierdo et al., 2017; Jackson et al., 2024) because vision is not the dominant sensory modality in rodents (Lankford et al., 2020). For instance, rats learn odour discrimination more quickly than visual discrimination in RLTs (Brushfield et al., 2008). For this reason, the bowl digging PRLT was predicted to show greater sensitivity to changes in cognitive flexibility. The lack of effects observed in either task may reflect differences in the behavioural mechanisms underlying performance in the human versus rat tasks. Cognitive flexibility in humans is strongly linked to the highly developed prefrontal cortex which may not be directly related to the neural circuits modulating apparently similar behaviours in rats.

In other behavioural models and cognitive flexibility assays, psilocybin has been shown to post-acutely improve reversal learning e.g. in a rodent model of anorexia (Conn et al., 2024). A subsequent study using the same paradigm in healthy rats showed that psilocybin reduced learning from negative feedback and increased learning from positive feedback (Fisher et al., 2024). These studies used a deterministic paradigm which differs from the PRLT in task difficulty. Furthermore, animals were testing using home cage feeding devices which allowed animals to voluntarily engage with the task, arguably providing a more naturalistic measure of motivation. In addition, they do not rely on food restriction which is known to alter physiology and brain function (Bubenik et al., 1992) and could have confounded flexibility behaviour in the current study. Both studies also used 1.5 mg/kg psilocybin which is higher than was used here however, the rat dose equivalent to the clinical trial dose of psilocybin is generally thought to align more closely with doses of 0.3-1 mg/kg (Higgins et al., 2021; Holze et al., 2023; Saito et al., 2004; Kolaczynska et al., 2021).

The variability in psychedelic effects on cognitive flexibility across animal and human studies underscores how the term *cognitive flexibility* is used inconsistently across the literature (Dajani & Uddin, 2017), encompassing a broad range of behavioural and neurocognitive processes. While the most frequently used proxy of cognitive flexibility is set-shifting, there is disagreement regarding this equivalency, with distinct research traditions emphasising different capabilities (e.g. generating multiple categorisations in creativity tasks or flexibility within language, mathematics, and perception) (Ionescu et al, 2024). In fact, recent conceptualisations distinguish *cognitive flexibility* (switching between different task sets according to changing rules) from *behavioural flexibility* (adjusting behaviour according to changing reward contingencies), Under this framework, while set-shifting tasks may be considered a measure of cognitive flexibility, reversal learning tasks such as the PRLT might be better conceptualised as measures of behavioural flexibility (Uddin, 2021). Accordingly, inconsistencies in the effects of psychedelics on flexibility may partly reflect differences in the specific aspect of flexibility being assessed.

## Conclusion

Overall, acute psilocybin treatment had no specific effects on rule learning in a touchscreen PRLT but induced dose-dependent impairments in task engagement and performance. The present findings are therefore more consistent with a general slowing of learning and non-specific behavioural disruption, possibly due to psilocybin’s perceptual effects. Any acute effects on learning were not sustained. Similarly, LSD produced only limited and inconsistent effects on PRLT behaviour, with scattered changes across doses and timepoints rather than a clear or robust influence on cognitive flexibility. Moreover, post-acute effects of psilocybin were not observed in a more ethologically valid bowl-digging PRLT variant, which was expected to be more sensitive to drug-induced changes in cognitive flexibility. Ultimately, these findings challenge the translational relevance of these tasks for assessing psilocybin effects on cognitive flexibility, with home cage variants potentially offering more sensitive and translationally relevant alternatives.

## Statements and Declarations

### Author Contributions

Dasha Anderson, Gary Gilmour, Chris Thomas, Caroline Golden and Emma SJ Robinson conceptualised the work. Dasha Anderson performed data collection of operant PRLT experiments and data analysis for all PRLT experiments. Nanni Maillot performed data collection for bowl digging PRLT experiments under the supervision of Dasha Anderson. Dasha Anderson prepared the original draft. All authors contributed to experimental design and planning as well as reviewing, editing and approval of the manuscript.

### Ethics Approval

All studies were performed in accordance with with the UK Animal (Scientific Procedures) Act, with ethical approval from the University of Bristol AWERB and under a Home Office Project license.

### Statement on Welfare of Animals

The sample size was based on previous rodent PRLT studies which we expected elicited a similar effect size to the current study (Jackson et al., 2024; Wilkinson et al., 2020). Animals were fully habituated to handling before initiating experiments and housed in enriched housing containing a red shelf, wooden chews, rope, and cardboard tube. Rats were provided with access to a playpen for up to 30 min per day during weeks when they were not training or testing (www.3hs-initiative.co.uk). Rats were maintained at no less than 90% their free-feeding weight and underwent weekly weighing and body scoring. Refined low-restraint intraperitoneal dosing methods were used during drug administration (www.3hs-initiative.co.uk).

## Data Availability

Raw data, Prism files, SPSS files, and code are available at: https://osf.io/e9dkg/overview?view_only=519e53607fc24f0ebc4b8615264cd348

## Acknowledgements

This work was supported by a BBSRC South West Bio doctoral studentship (BB/T008741/1) and BBSRC project grant (BB/V015028/1) in collaboration with Compass Pathways plc.

