## Supplementary material for "Investigating the Effects of Psilocybin on Cognitive Flexibility in Touchscreen and Naturalistic Variations of the Probabilistic Reversal Learning Task"

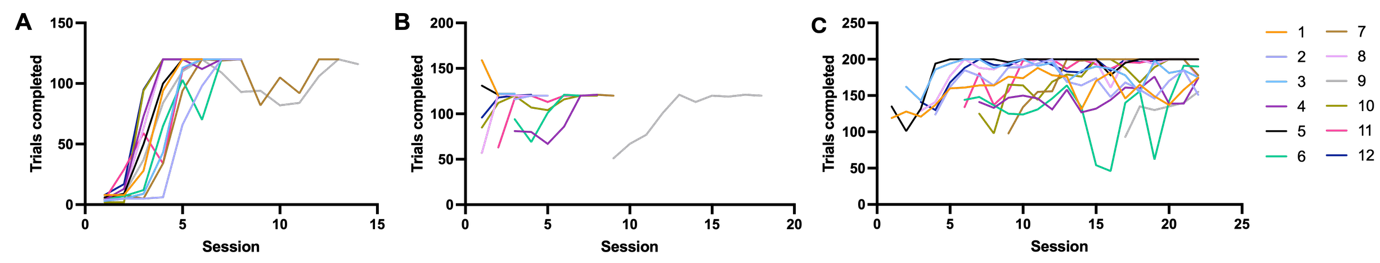

**Supplementary Fig S1** Training data across CRF1, CRF2, and the PRLT. *(****A****) Number of trials completed in CRF1. (****B****) Number of trials completed in CRF2. (****C****) Number of trials completed in the PRLT*

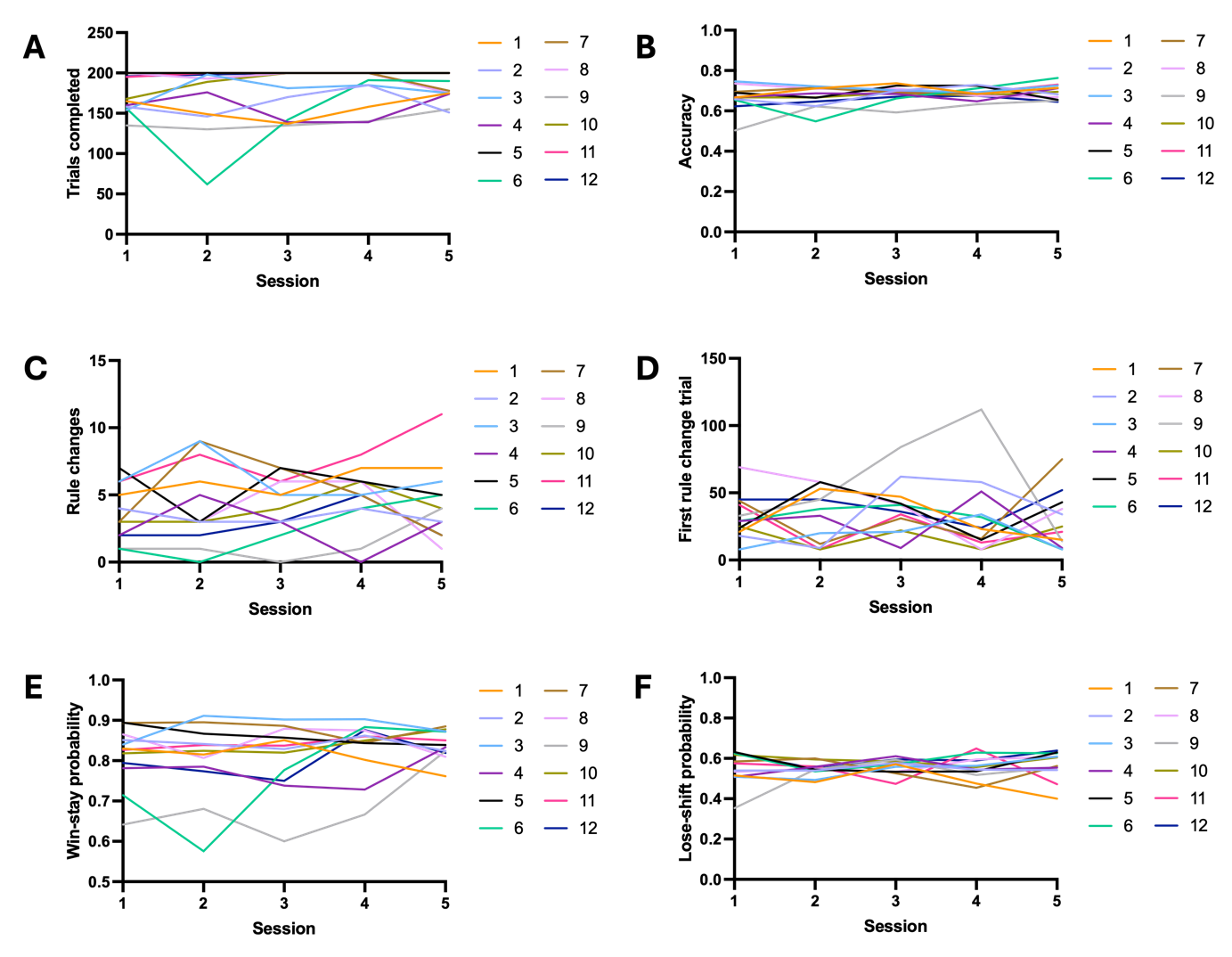

**Supplementary Fig S2** Stability of animals’ performance in the final 5 sessions of training. *(****A****) Number of trials completed in a session. (****B****) Accuracy (percentage of total trials that were correct responses). (****C****) Number of rule changes completed in a session. (****E****) Trial at which animals achieved their first rule change in a session. (****F****) Win stay probability (likelihood of selecting the previously rewarded stimulus). (****G****) Lose shift probability (likelihood of avoiding the previously rewarded stimulus)*

**Supplementary Methods S3. Computational modelling**

The first (Q-learn1) was a model adapted from (Grogan et al., 2017) based on the Q-learning model, which assumes that the animals learned the task according to reward prediction error (RPE) (Rescorla, 1972). For each stimulus *i* it is assumed that the animal estimates the reward value associated with that stimulus, *Q*(*i*). Initially this value was set at 0.5 and updated with each trial after feedback is received according to:

$$Q_{t+1}\left( i \right)=Q_{t}\left( i \right)+\alpha\delta$$

where *α* is the learning rate and δ is the RPE, the difference between the reward received and the expected reward:

$$\delta=r-Q_{t}(i)$$

In the second model (Q-learn2), expected values were updated based on a learning rate *α* that was separate for positive and negative RPEs:

if *δ*=1

if *δ*=0

$$Q_{t+1}(i)= \left\{ \begin{aligned} Q_{t}+\alpha_{+}\delta\\ Q_{t}+\alpha\_\delta\end{aligned} \right.$$

The probability of choosing each stimulus was calculated using the softmax function:

$$P(i)=\frac{e^{\beta Q_{t}(i)}}{\sum_{i}^{n} e^{{\beta Q}_{t}(i)}}$$

where *β* is the inverse temperature of the softmax equation, a free parameter which determines the extent to which the Q value of different stimuli affects choice probabilities, with high *β* values meaning they have a large effect (choices are more ‘deterministic’) and low *β* values meaning they have a smaller effect (choices are more ‘random’) (Nussenbaum, 2019). For each animal, both models were fit to the PRLT and the best parameters for each animal were identified as those associated with the lowest sum of the negative log-likelihood (*nll*). Learning rates and *β* values were randomly initialised between 0.01 and 0.99 and then optimised.

The best-fitting model was then identified using the Bayesian Information Criterion (BIC) (Schwarz, 1978):

$$\left( \log(n) \right) k-2ll$$

where *n* is the number of trials, *k* is the number of parameters in the model and *ll* is the log-likelihood of the model. The model with the lowest BIC score was deemed the best-fitting model.

**Supplementary Methods S4. Foraging training**

On the first day of habituation, rats explored the arena for 10 minutes with their cage mate and on the second day they explored the arena alone for 10 minutes. In the following session, the bowls were baited with a reward pellet at either 10 o’clock, 12 o’clock, or 2 o’clock positions and the rat was allowed to explore the bowl to find the pellet. The front left corner of the arena was then baited to encourage the animal to return to the start point. Bowls were rebaited for the next trial, and this was repeated for 12 trials. If animals failed to collect the pellet after 30 seconds, they were removed from the arena for 5 seconds before re-starting the trial.

In the subsequent three sessions rats underwent digging training. The pellet was placed in the same location but covered with 1 cm of sawdust, while the other was left empty. This was then repeated for another 12 trials, except rats were given 15 seconds to find the pellet, otherwise it was removed for 5 seconds and scored as an omission. The position of the baited bowl was pseudo-randomised each trial. The same procedure was used the next two days of training, except a 2 cm layer of sawdust was used instead of 1 cm and the non-baited bowl was removed once the animal found the pellet. By day 3, the front left corner was no longer rebaited.

After digging training, animals underwent three discrimination tests in which each bowl was filled with 2cm of digging substrate (e.g. cardboard squares, sponge, shredded cloth, etc). The baited substrate was counterbalanced across animals. Finely crushed pellet was mixed into both substrates to prevent animals using olfactory cues to locate the pellets. One of the substrates was consistently baited and the position of bowls was pseudo-randomised. Animals were left to explore the bowls to find the pellet, with the non-chosen bowl removed from the second trial. In all sessions, every animal achieved the 6-trial criterion within 20 trials. Once the animal correctly chose the baited bowl on six consecutive trials within 20 trials, the animal was considered fully trained and ready for probabilistic learning sessions.

Rats underwent one probabilistic learning session where one substrate was baited with a single pellet 80% of the time (the ‘rich’ substrate) and the other 20% of the time (the ‘lean’ substrate). As before, the position of the bowls was pseudo-randomised across trials and the rich and lean substrate were counterbalanced across animals. The rat had to choose the correct (rich) bowl 6 consecutive times within a maximum of 30 trials to successfully acquire the first rule. If the animal did not make a choice within 30 seconds, they were removed from the arena for 10 seconds and this was recorded as an omission. The session ended after either 30 minutes, five consecutive omissions, successful acquisition of the first rule, or after failing to acquire the rule within 30 trials. Eight animals acquired the rule within 30 trials. Ten animals reversed in the baseline PRLT session, with only two animals acquiring the first rule and not reversing. The average number of trials to reach the first rule acquisition was 23.1±11.8. Amongst those that reversed, the average number of reversals was 1.8±0.871. No animals failed to learn the first rule within 50 trials.

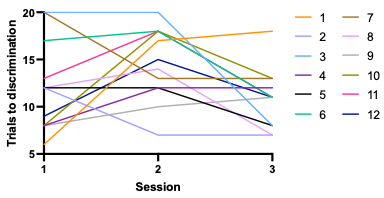

***Supplementary Fig S5*** Number of trials to reach criterion across three discrimination sessions. *Animals were required to achieve six consecutive correct trials in order to reach the criterion for discrimination*

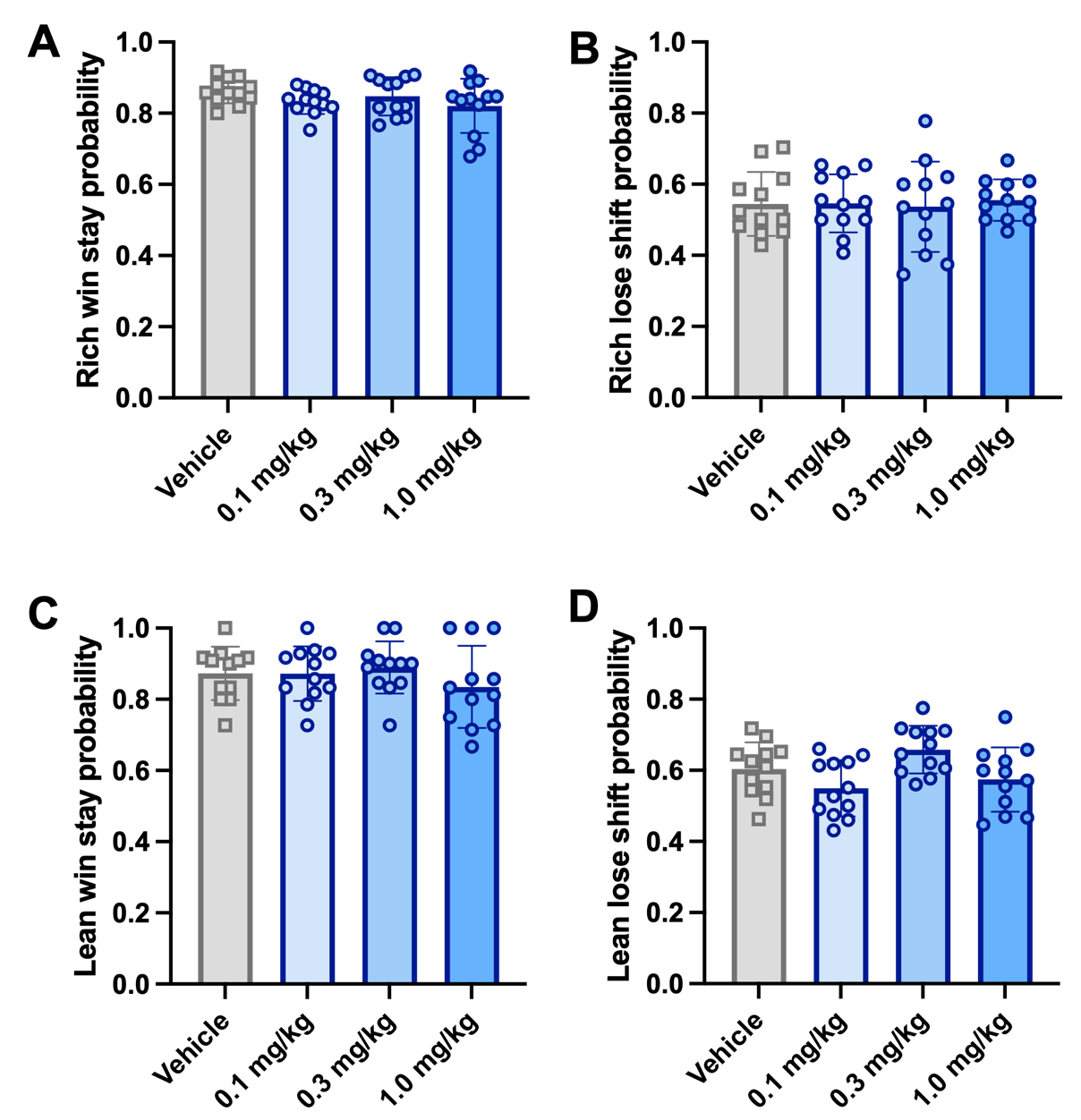

***Supplementary Fig S6*** Acute effects of psilocybin on win stay and lose shift probability in the touchscreen PRLT when split by rich and lean stimulus. *(****A****) Rich win stay probability (likelihood of selecting the previously rewarded stimulus when it is more frequently rewarded overall). (****B****) Rich lose shift probability (likelihood of selecting the previously rewarded stimulus when it is less frequently rewarded overall). (****C****) Lean win stay probability (likelihood of avoiding the previously unrewarded stimulus when it is more frequently rewarded overall). (****D****) Lean lose shift probability (likelihood of avoiding the previously unrewarded stimulus when it is less frequently rewarded overall). Data shown as mean +/- sem and individual data points represent individual subjects, n = 12, *p<0.05, **p<0.01, ***p<0.001 post-hoc pairwise comparison vs vehicle control*

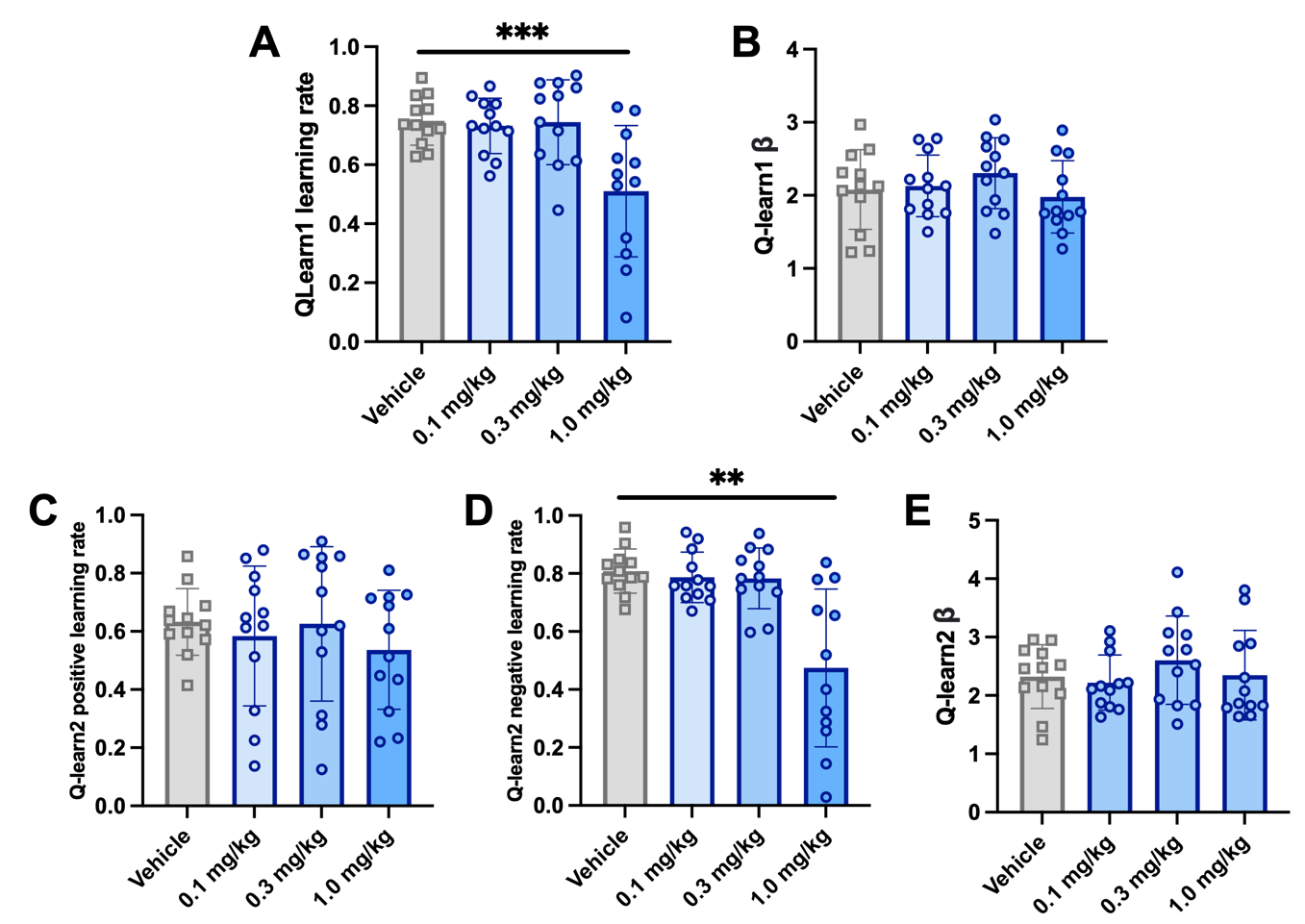

***Supplementary Fig S7*** Acute effects of psilocybin on Q-learning model outputs in the touchscreen PRLT. *(****A****) Q-learn1 (single learning rate model) learning rate (α), a parameter which determines to what extent the current Q-value is updated based on the reward prediction error. (****B****) Q-learn1 (single learning rate model) β, a parameter reflecting choice determinism. (****C****) Q-learn2 (dual learning rate model) learning rate (α). (****D****) Q-learn2 (dual learning rate model) negative learning rate (α). (****E****) Q-learn2 (dual learning rate model) β. Data shown as mean +/- sem and individual data points represent individual subjects, n = 12, *p<0.05, **p<0.01, ***p<0.001 post-hoc pairwise comparison vs vehicle control*

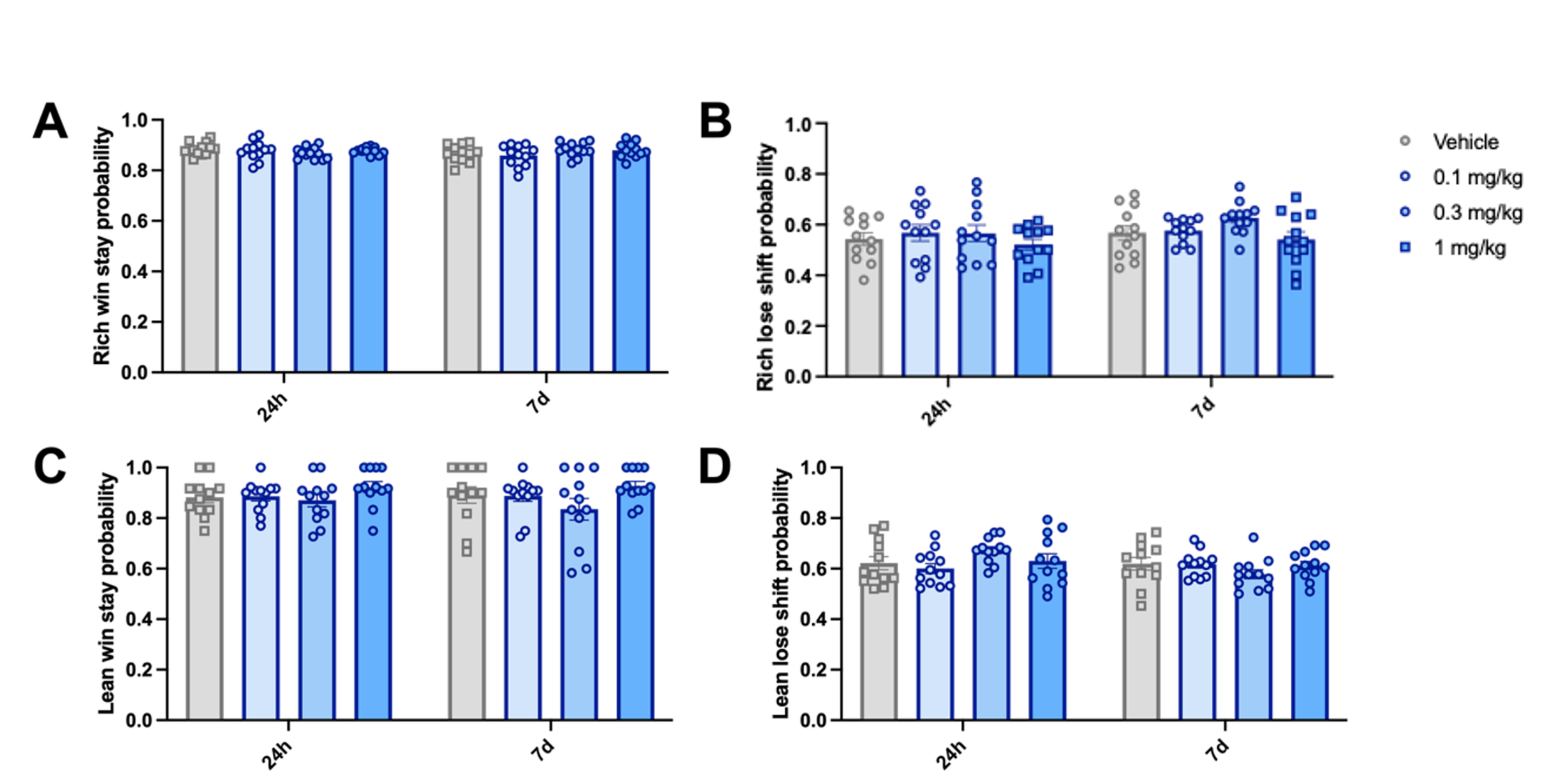

***Supplementary Fig S8*** Post-acute effects of psilocybin on win stay and lose shift probability in the touchscreen PRLT when split by rich and lean stimulus at 24 hours and 7 days post-dose. *(****A****) Rich win stay probability (likelihood of selecting the previously rewarded stimulus when it is more frequently rewarded overall). (****B****) Rich lose shift probability (likelihood of selecting the previously rewarded stimulus when it is less frequently rewarded overall). (****C****) Lean win stay probability (likelihood of avoiding the previously unrewarded stimulus when it is more frequently rewarded overall). (****D****) Lean lose shift probability (likelihood of avoiding the previously unrewarded stimulus when it is less frequently rewarded overall). Data shown as mean +/- sem and individual data points represent individual subjects, n = 12, *p<0.05, **p<0.01, ***p<0.001 post-hoc pairwise comparison vs vehicle control*

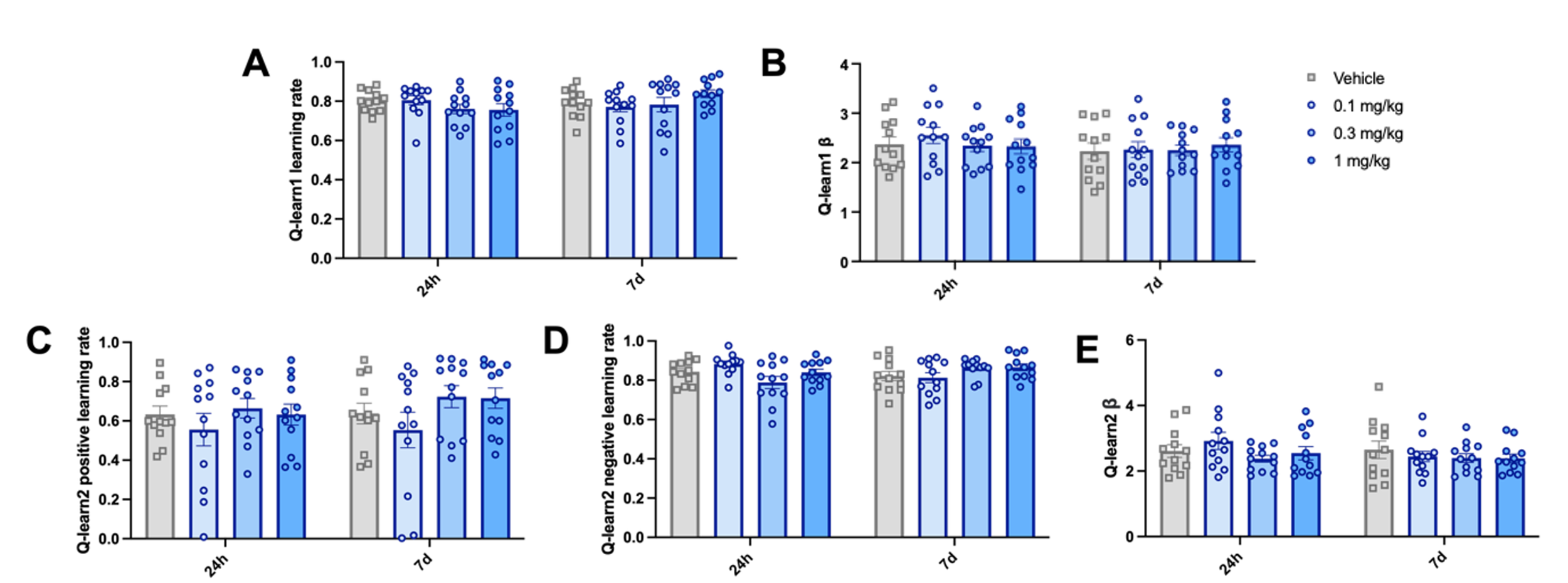

***Supplementary Fig S9*** Post-acute effects of psilocybin on Q-learning model outputs in the touchscreen PRLT at 24 hours and 7 days post-dose. *(****A****) Q-learn1 (single learning rate model) learning rate (α), a parameter which determines to what extent the current Q-value is updated based on the reward prediction error. (****B****) Q-learn1 (single learning rate model) β, a parameter reflecting choice determinism. (****C****) Q-learn2 (dual learning rate model) learning rate (α). (****D****) Q-learn2 (dual learning rate model) negative learning rate (α). (****E****) Q-learn2 (dual learning rate model) β. Data shown as mean +/- sem and individual data points represent individual subjects, n = 12, *p<0.05, **p<0.01, ***p<0.001 post-hoc pairwise comparison vs vehicle control*

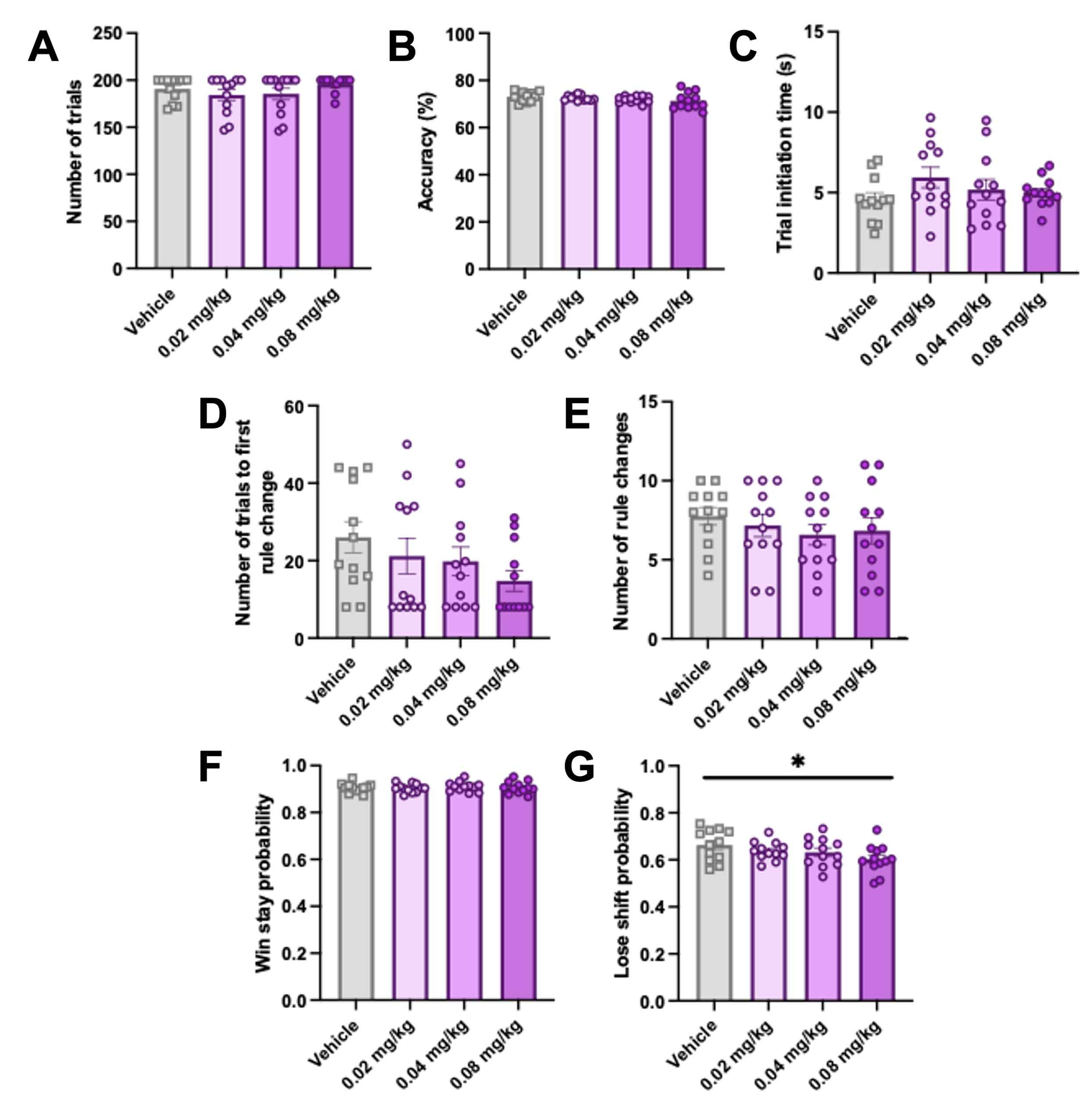

***Supplementary Fig S10*** Acute effects of LSD on main operant PRLT output measures. *(****A****) Number of trials completed in a session. (****B****) Accuracy (percentage of total trials that were correct responses). (****C****) Time taken for animals to self-initiate each trial. (****D****) Number of rule changes completed in a session. (****E****) Trial at which animals achieved their first rule change in a session. (****F****) Win stay probability (likelihood of selecting the previously rewarded stimulus). (****G****) Lose shift probability (likelihood of avoiding the previously rewarded stimulus). Data shown as mean +/- sem and individual data points represent individual subjects, n = 11, *p<0.05, **p<0.01, ***p<0.001 post-hoc pairwise comparison vs vehicle control*

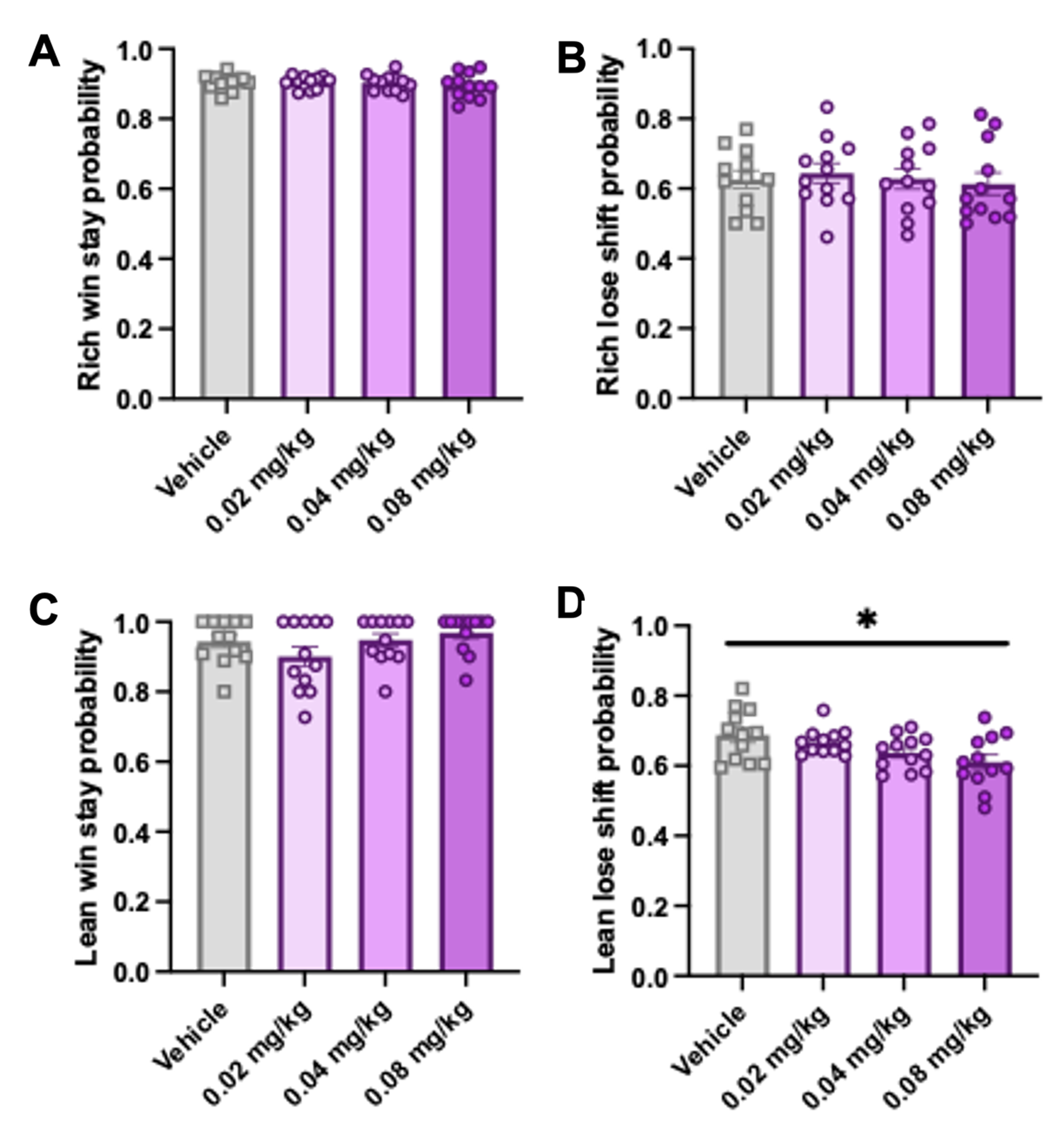

***Supplementary Fig S11*** Acute effects of LSD on win stay and lose shift probability in the touchscreen PRLT when split by rich and lean stimulus. *(****A****) Rich win stay probability (likelihood of selecting the previously rewarded stimulus when it is more frequently rewarded overall). (****B****) Rich lose shift probability (likelihood of selecting the previously rewarded stimulus when it is less frequently rewarded overall). (****C****) Lean win stay probability (likelihood of avoiding the previously unrewarded stimulus when it is more frequently rewarded overall). (****D****) Lean lose shift probability (likelihood of avoiding the previously unrewarded stimulus when it is less frequently rewarded overall). Data shown as mean +/- sem and individual data points represent individual subjects, n = 11, *p<0.05, **p<0.01, ***p<0.001 post-hoc pairwise comparison vs vehicle control*

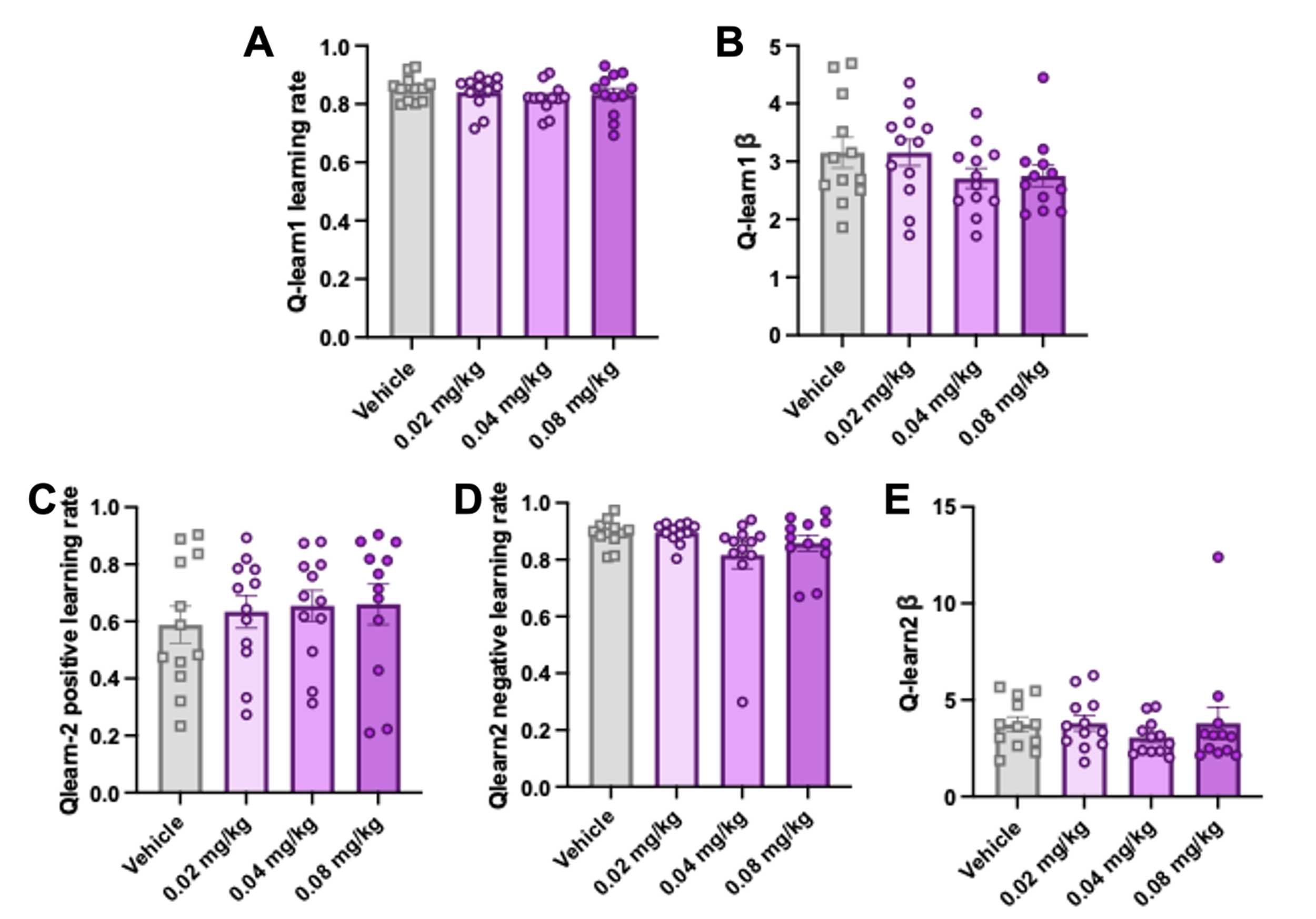

***Supplementary Fig S12*** Acute effects of LSD on Q-learning model outputs in the touchscreen PRLT. *(****A****) Q-learn1 (single learning rate model) learning rate (α), a parameter which determines to what extent the current Q-value is updated based on the reward prediction error. (****B****) Q-learn1 (single learning rate model) β, a parameter reflecting choice determinism. (****C****) Q-learn2 (dual learning rate model) learning rate (α). (****D****) Q-learn2 (dual learning rate model) negative learning rate (α). (****E****) Q-learn2 (dual learning rate model) β. Data shown as mean +/- sem and individual data points represent individual subjects, n = 11, *p<0.05, **p<0.01, ***p<0.001 post-hoc pairwise comparison vs vehicle control*

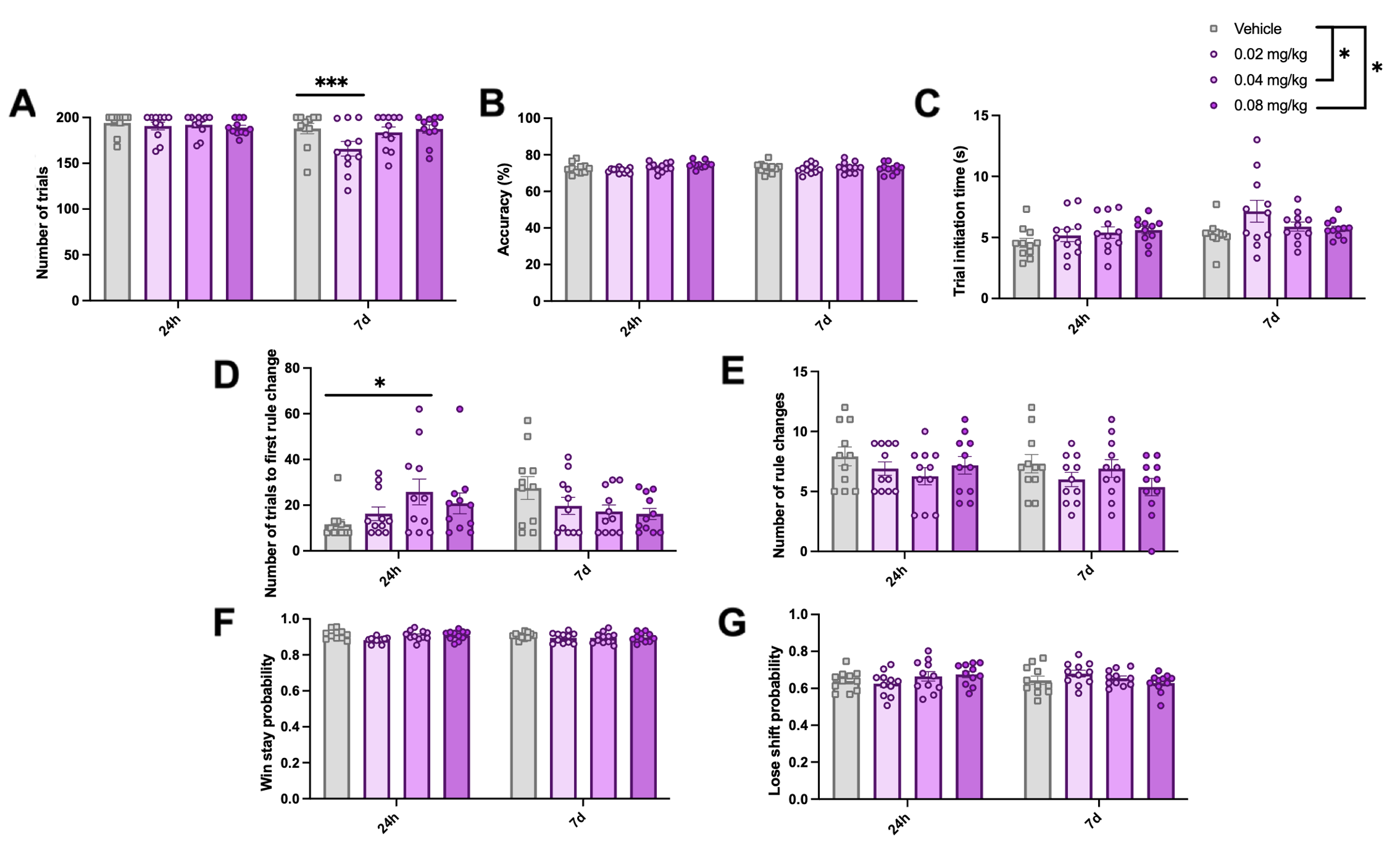

***Supplementary Fig S13*** Post-acute effects of LSD on main operant PRLT output measures at 24 hours and 7 days post-dose. *(****A****) Number of trials completed in a session. (****B****) Accuracy (percentage of total trials that were correct responses). (****C****) Time taken for animals to self-initiate each trial. (****D****) Number of rule changes completed in a session. (****E****) Trial at which animals achieved their first rule change in a session. (****F****) Win stay probability (likelihood of selecting the previously rewarded stimulus). (****G****) Lose shift probability (likelihood of avoiding the previously rewarded stimulus). Data shown as mean +/- sem and individual data points represent individual subjects, n = 11, *p<0.05, **p<0.01, ***p<0.001 post-hoc pairwise comparison vs vehicle control*

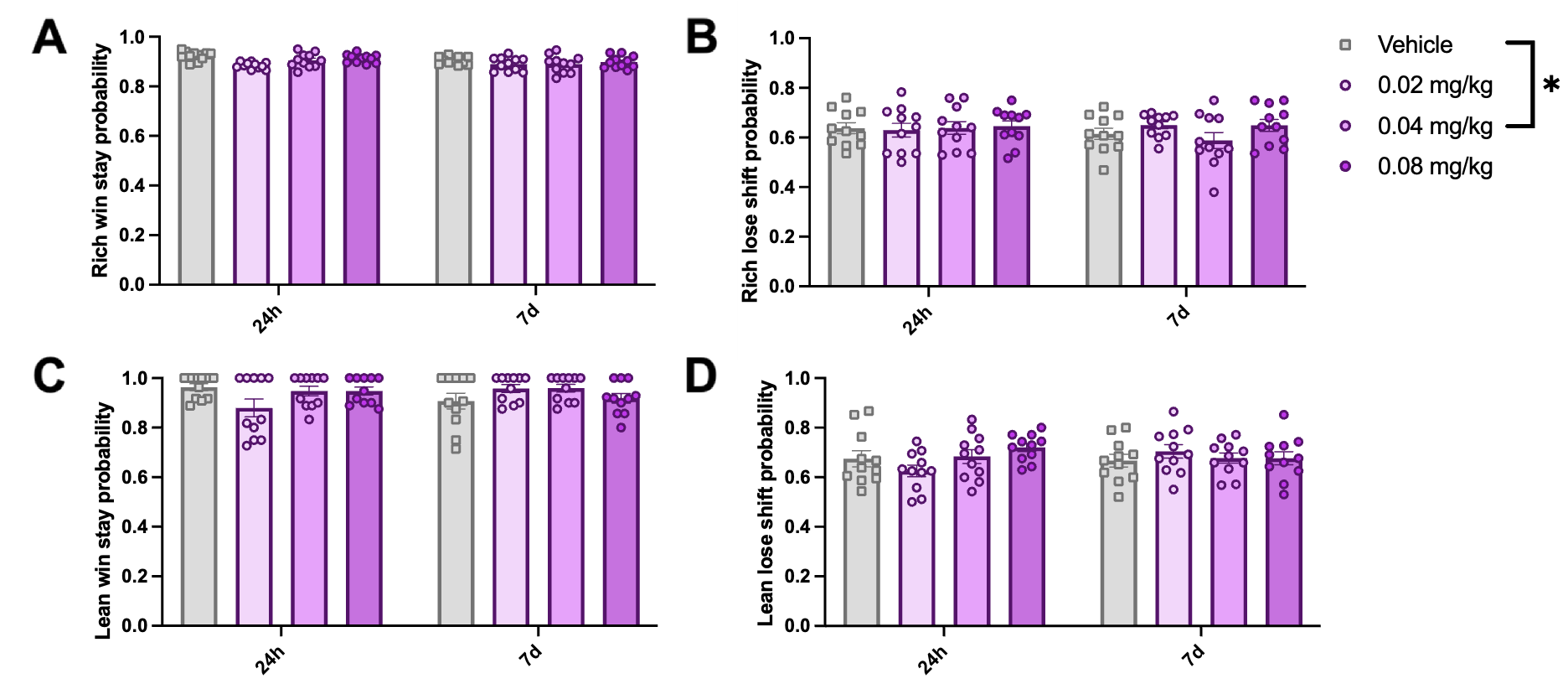

***Supplementary Fig S14*** Post-acute effects of LSD on win stay and lose shift probability in the touchscreen PRLT when split by rich and lean stimulus at 24 hours and 7 days post-dose. *(****A****) Rich win stay probability (likelihood of selecting the previously rewarded stimulus when it is more frequently rewarded overall). (****B****) Rich lose shift probability (likelihood of selecting the previously rewarded stimulus when it is less frequently rewarded overall). (****C****) Lean win stay probability (likelihood of avoiding the previously unrewarded stimulus when it is more frequently rewarded overall). (****D****) Lean lose shift probability (likelihood of avoiding the previously unrewarded stimulus when it is less frequently rewarded overall). Data shown as mean +/- sem and individual data points represent individual subjects, n = 11, *p<0.05, **p<0.01, ***p<0.001 post-hoc pairwise comparison vs vehicle control.*

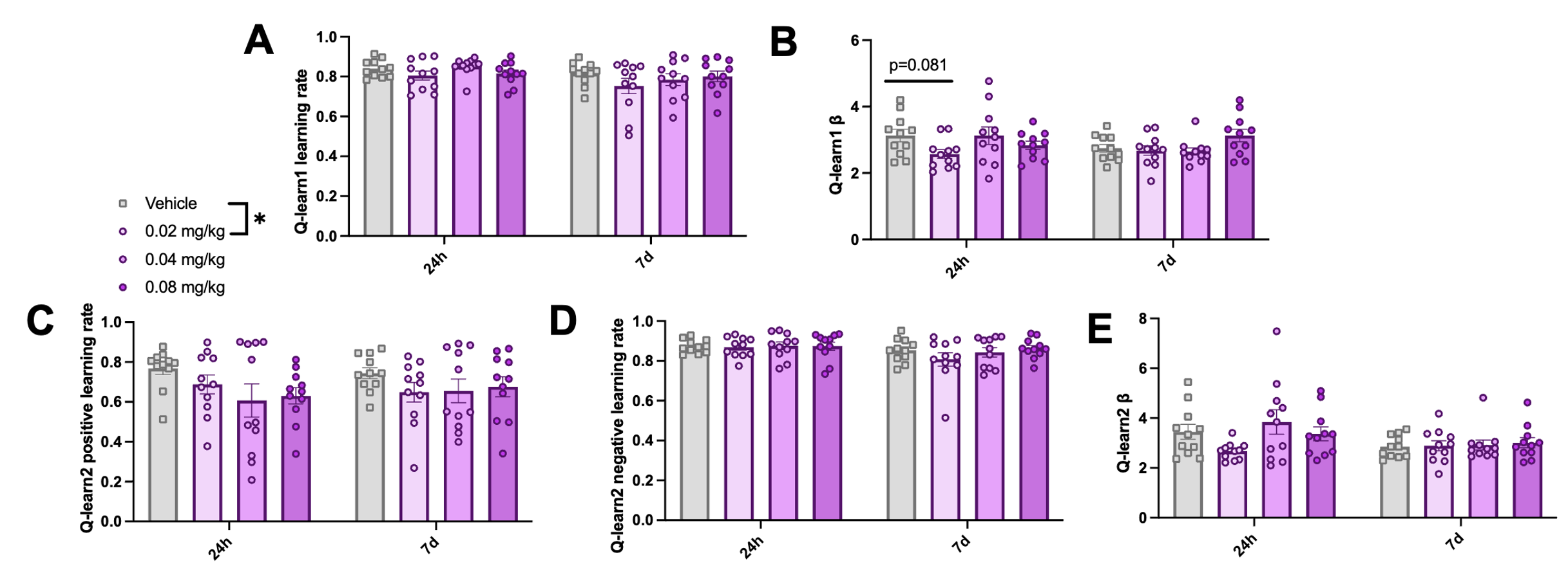

***Supplementary Fig S15*** Post-acute effects of LSD on Q-learning model outputs in the touchscreen PRLT at 24 hours and 7 days post-dose. *(****A****) Q-learn1 (single learning rate model) learning rate (α), a parameter which determines to what extent the current Q-value is updated based on the reward prediction error. (****B****) Q-learn1 (single learning rate model) β, a parameter reflecting choice determinism. (****C****) Q-learn2 (dual learning rate model) learning rate (α). (****D****) Q-learn2 (dual learning rate model) negative learning rate (α). (****E****) Q-learn2 (dual learning rate model) β. Data shown as mean +/- sem and individual data points represent individual subjects, n = 11, *p<0.05, **p<0.01, ***p<0.001 post-hoc pairwise comparison vs vehicle control*

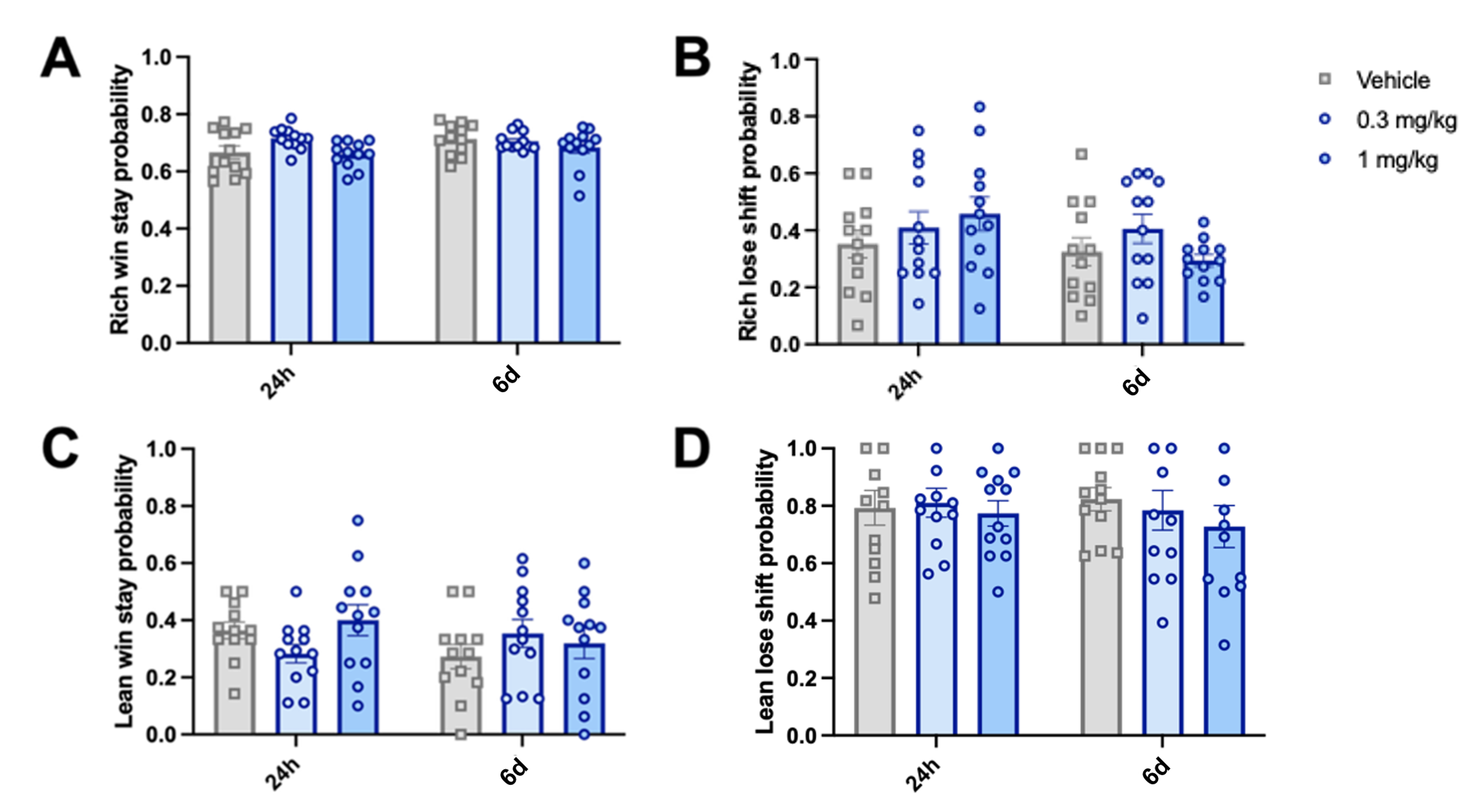

***Supplementary Fig S16*** Post-acute effects of psilocybin on win stay and lose shift probability in the foraging PRLT when split by rich and lean stimulus at 24 hours and 6 days post-dose*. (****A****) Rich win stay probability (likelihood of selecting the previously rewarded stimulus when it is more frequently rewarded overall). (****B****) Rich lose shift probability (likelihood of selecting the previously rewarded stimulus when it is less frequently rewarded overall). (****C****) Lean win stay probability (likelihood of avoiding the previously unrewarded stimulus when it is more frequently rewarded overall). (****D****) Lean lose shift probability (likelihood of avoiding the previously unrewarded stimulus when it is less frequently rewarded overall). Data shown as mean +/- sem and individual data points represent individual subjects, n = 12, *p<0.05, **p<0.01, ***p<0.001 post-hoc pairwise comparison vs vehicle control*
